# Proteomic identification and validation of novel neuronal EV-based markers for Alzheimer’s disease biomarker discovery

**DOI:** 10.64898/2026.04.26.720870

**Authors:** Waqar Ahmed, Carlos Nogueras-Ortiz, Ram Sagar, Daiyun Dong, Rachel J Boyd, Pamela J Yao, Anton Iliuk, Constantine G. Lyketsos, Kenneth W. Witwer, Dimitrios Kapogiannis, Vasiliki Mahairaki

## Abstract

Extracellular vesicles (EVs) circulate in biofluids and carry tissue-specific molecular cargo, offering significant potential for the discovery of minimally invasive biomarkers. However, translation in neurodegenerative diseases has been hindered by the lack of validated neuronal EV surface markers that enable selective isolation from plasma. We hypothesized that proteomic profiling of EVs released from human induced pluripotent stem cell (hiPSC)-derived neurons would identify 1. robust Alzheimer’s disease (AD)-associated signatures that reflect disease pathogenesis, and 2. surface-accessible neuronal markers capable of enriching disease-relevant cargo. Neurons differentiated from AD patients and age-matched cognitively normal (CN) individuals were used to isolate EVs, which were characterized and analyzed by LC-MS proteomics in both total and membrane-enriched fractions. Proteomic profiling identified numerous dysregulated proteins, with a subset validated across independent AD datasets. We identified CNTNAP2 and STX1B as neuronal, brain-enriched EV surface proteins accessible for selective capture and confirmed their presence in EVs from post-mortem human brain, supporting them as bona fide brain-derived EV markers. Immuno-isolation of plasma EVs showed that CNTNAP2-positive EVs had a robust AD-associated increase in phosphorylated tau, identifying CNTNAP2 as a highly discriminative brain-derived EV marker and supporting its potential for blood-based AD diagnostics.

**Graphical Abstract:** Previous studies indicate that extracellular vesicles (EVs) released from neurons carry disease-relevant cargo, yet the search for the optimal neuronal surface markers for the selective isolation of EVs pertinent to Alzheimer’s disease (AD) from blood is ongoing. To address this need, we performed proteomic profiling of EVs derived from hiPSC-neurons (iNEVs) of AD patients and cognitively normal individuals (CN). LC-MS analysis of whole and membrane-enriched EV fractions revealed robust protein dysregulation and identified CNTNAP2 and STX1B as surface-exposed neuronal EV proteins, confirmed in human brain-derived EVs (BDEVs). Immuno-isolation of plasma EVs demonstrated that CNTNAP2-positive EVs are enriched for phosphorylated tau in AD, whereas the previously used markers; NrCAM and ATP1A3-positive EVs showed limited discrimination. These findings establish a robust framework for proteomic discovery and nominate CNTNAP2 as a promising EV selection marker for blood-based AD diagnostics.

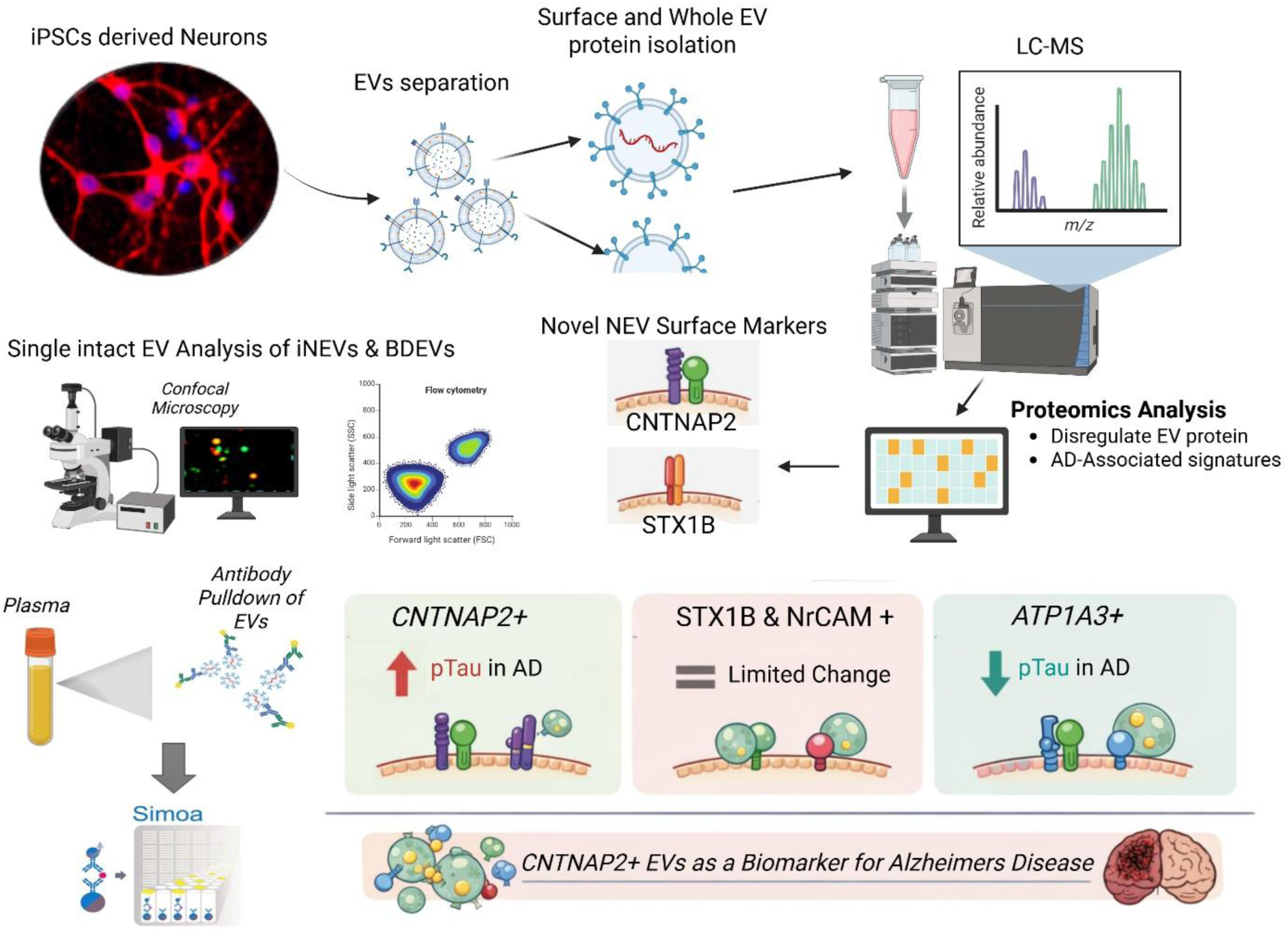

## 1 INTRODUCTION

Extracellular vesicles (EVs) are nano-sized membranous particles that are continuously being released by cells into the extracellular environment. These vesicles play crucial roles in intercellular communication and have potential for biomarker discovery and therapeutic applications. EVs originate from various tissues throughout the body and circulate in blood plasma, making them a valuable source of information about the physiological and pathological states of different organs (Pulliam et al. 2019; Bahmani and Ullah 2022; Yuana et al. 2013) that can be acquired through a simple blood draw. For neurodegenerative diseases, such as Alzheimer’s and Parkinson’s disease, early detection and monitoring are critical for effective management and intervention. The isolation of neuronal EVs (NEVs) in human plasma (Campbell and Mocchetti 2021; Coleman and Hill 2015; Manu et al. 2021; Upadhya and Shetty 2021) offer a unique opportunity to access brain-specific biomarkers that could aid in early detection or diagnosis and have the potential to extend the window of therapeutic intervention. However, reliably isolating organ specific EVs remains a challenge. Studies over the last decade have identified EV surface antigens that permit the isolation of neuron-specific EVs. For example, L1 cell adhesion molecule (L1CAM/CD171) and Neural Cell Adhesion Molecule 1 (NCAM1/CD56) were originally identified as surface markers that permit the immunoaffinity capture of NEVs from plasma (Fiandaca et al. 2015). In the last decade, L1CAM+ EVs have since been used to isolate NEVs from plasma in a variety of studies (Mustapic et al. 2017; Gill et al. 2018; Markaki et al. 2023; Liao et al. 2025; Qin et al. 2022) and L1CAM was validated as an NEV surface protein at the single EV level (Pulliam et al. 2019). However, according to The Human Protein Atlas (https://www.proteinatlas.org/), while L1CAM and NCAM1 are highly enriched in brain tissues, their expression is not exclusively restricted to the brain (Hill 2019). Moreover, L1CAM is upregulated during cancer progression(D. E. Gomes and Witwer 2022). Therefore, there is a need to identify neuronal-specific markers for isolating more pure populations of NEVs.

Proteomics has emerged as a powerful tool for characterizing the protein composition of EVs. You *et al*. used such an approach to identify cell type-specific EV protein markers for excitatory neurons, astrocytes, microglia-like cells, and oligodendrocyte-like cells (You et al. 2023, 2022). In addition to identifying 16 common proteins shared across EVs from all studied cell types in the brain, they reported that ATPase Na+/K+ transporting subunit alpha 3 (ATP1A3) and NCAM1 are abundantly expressed in EVs isolated from human iPSC-derived neurons (You et al. 2023, 2022). However, data from The Human Protein Atlas also suggests that ATP1A3 is highly enriched in heart muscle; thus, reducing confidence in the specificity of this marker for isolating NEVs. By focusing the analysis on the surface proteome of EVs, specific markers can be identified to facilitate the isolation of EVs derived from distinct tissues.

In this study, we performed surface and whole EVs proteomics and bioinformatic analysis to identify additional EV surface proteins that are exclusively enriched in the brain. We utilized random forest modeling to identify AD-associated biomarkers that can be used to discriminate AD-NEVs from NEVs isolated from cognitively normal individuals (CN). We further validated the newly identified neuronal EV surface markers by isolating different populations of NEVs from plasma and brain derived EVs (BDEVs) from individuals with AD and CN, revealing selection marker-dependent differences in disease-associated cargo. The newly identified markers outperformed previously used neuronal markers in discriminating between AD patients and CN, underscoring their potential as disease biomarkers. Our study provides a framework that can be used to enhance blood biomarker development for brain diseases, potentially improving early detection and patient outcomes.

## 2 MATERIALS & METHODS

### 2.1 Study Participants

#### Collection of Human Peripheral Blood Mononuclear Cells (PBMCs)

We obtained peripheral blood mononuclear cells (PBMCs) derived from blood samples by the Genetic Core Facility at Johns Hopkins. Blood samples were obtained from clinically well-characterized participants in the Johns Hopkins Alzheimer’s Disease Research Center (JHADRC) cohort. The study included fifteen participants, with 8 having AD and 7 serving as CN controls; demographic characteristics such as age, gender, and *APOE* genotype are in Table 1. Diagnoses of AD or CN were established by an JHADRC clinical diagnostic panel of experts. All participants or their legal representatives provided informed consent under the oversight of a Johns Hopkins Institutional Review Board (NA_00045104) prior to blood sampling.

**Table 1.**
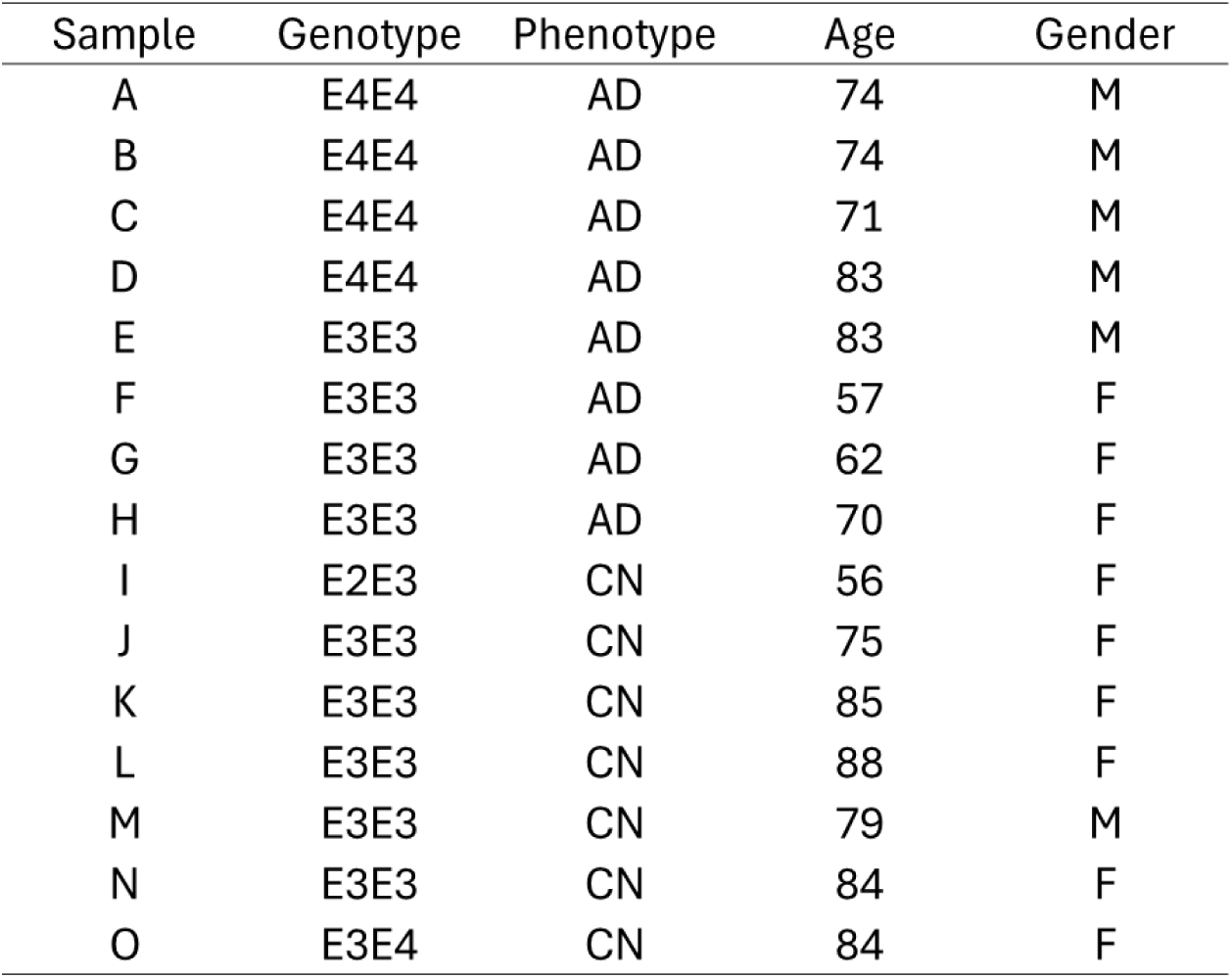
Clinical and demographic information of n = 15 study participants.

| Sample | Genotype | Phenotype | Age | Gender |
| --- | --- | --- | --- | --- |
| A | E4E4 | AD | 74 | M |
| B | E4E4 | AD | 74 | M |
| C | E4E4 | AD | 71 | M |
| D | E4E4 | AD | 83 | M |
| E | E3E3 | AD | 83 | M |
| F | E3E3 | AD | 57 | F |
| G | E3E3 | AD | 62 | F |
| H | E3E3 | AD | 70 | F |
| I | E2E3 | CN | 56 | F |
| J | E3E3 | CN | 75 | F |
| K | E3E3 | CN | 85 | F |
| L | E3E3 | CN | 88 | F |
| M | E3E3 | CN | 79 | M |
| N | E3E3 | CN | 84 | F |
| O | E3E4 | CN | 84 | F |

#### Human Brain Tissue Collection

We obtained unfixed, frozen postmortem human brain tissue from the Brain Resource Center (BRC) at Johns Hopkins University. The cohort consisted of 6 individuals diagnosed with AD and 6 CN controls; characteristics such as age, gender, CERAD and BRAAK scores are provided in Supplementary Table 1. Clinical diagnoses were confirmed according to established CERAD and BRAAK criteria, as previously described (Murayama and Saito 2004). Approximately 200 mg of tissue was collected from the superior and middle temporal gyrus corresponding to Brodmann areas 40, 42, and 21. The brain tissue was stored at −80°C until EV separation and characterization.

### 2.2 Induced pluripotent stem cell culture and neuronal differentiation

We generated human induced pluripotent stem cells (hiPSCs) from PBMCs derived from study participants, following a previously established protocol (Zhang et al. 2013; Sagar et al. 2023). For neuronal differentiation, all iPSC lines were transduced with a lentivirus expressing NgN2 and rtTA, following a previously established protocol to generate a homogeneous population of excitatory neurons within two weeks (Sagar et al. 2023; Zhang et al. 2013). For induction of neuronal differentiation, NgN2-positive cells were seeded on vitronectin-coated plates at a density of 250,000 cells per well in Essential 8 (E8) medium supplemented with a ROCK Inhibitor Y-27632. Cells were maintained in E8 medium until 80% confluency, and doxycycline was added to induce NgN2 expression (Day 0). On Day 1, cells were selected using puromycin in the induction medium and passaged onto Matrigel-coated 6-well plates the following day. On Day 4, 50% of the media was replaced with neuronal maturation media supplemented with Ara-C to inhibit non-neuronal cell proliferation. From Day 6 to Day 12, 70% of the neuronal maturation media was changed every other day. From Day 15 to Day 50, 50% of the maturation media, without doxycycline, was changed every 72 hrs. On day 50 of mature neuronal cells condition medium was collected for EV separation and characterization.

### 2.3 Extracellular vesicle separation

At day 50 of neuronal differentiation EV were separated following a previously published protocol (Sagar et al. 2025). Briefly, 12 ml of cell supernatant from each neuronal culture was collected and centrifuged at 300 g for 10 minutes at 4°C. The resulting supernatant was transferred to a fresh Falcon tube and centrifuged at 2000 g for 10 minutes at 4°C to remove cell debris. The supernatant was transferred to a fresh tube and stored at -80°C. Supernatant was thawed and concentrated to 500 μl or less using an Amicon Ultra-15 centrifugal filter unit (Ultracel-100KD; 15 ml capacity; Merck Millipore, MA) by centrifugation at 2000 g at 4°C for 20 minutes. The concentrate was loaded onto a size exclusion chromatography column (qEV-70; IZON Science, Cambridge, MA) and eluted with DPBS using an automated fraction collector (AFC: IZON, Cambridge, MA). EV-containing fractions totaling 2 ml were collected after discarding the void volume (3.8 ml) and further concentrated using an Amicon Ultra-2 centrifugal filter unit (Ultracel-10KD; Merck Millipore, MA) to a final volume of 200 μl. EV preparations from AD patients (ADiNEVs) and CN (iNEVs) were placed into protein LoBind tubes (Cat# 13-698-794, Eppendorf) and stored at -80°C until further analysis.

### 2.4 Brain extracellular vesicle separation

Unfixed frozen cerebral cortex tissue (Brodmann areas 40, 42 or 21) from postmortem human brains was obtained from the Johns Hopkins Department of Pathology. Brain-derived EVs were isolated using a standardized workflow as previously described (Huang et al. 2023). Briefly, tissue was enzymatically dissociated, quenched with protease and phosphatase inhibitors, and cleared by sequential low-speed centrifugation and filtration. Large and small EV fractions were separated by differential centrifugation, size exclusion chromatography, and ultracentrifugation. EV pellets were resuspended in inhibitor-containing PBS, aliquoted, and stored at −80°C until analysis.

### 2.5 Nano-flow cytometry measurements (NanoFCM)

The particle number concentration and size distribution of EV preparations were determined using the Flow NanoAnalyzer (NanoFCM) following the manufacturer’s guidelines and established protocols (Arab et al. 2021; Huang et al. 2023). The instrument was calibrated with 250 nm fluorescently labelled silica beads and a NanoFCM™ Silica Nanospheres Cocktail #1 (Cat# S16M-Exo) for concentration and size distribution, respectively. EV samples were serially diluted in DPBS (typically from 1:100 to 1:200), and events were recorded for 1 minute. Particle numbers were calculated using a calibration curve based on flow rate and side scatter intensity.

### 2.6 Transmission Electron Microscopy (TEM)

For ultrastructural imaging, 10 μL of EV suspension (at approx. 4.0 × 10⁸ particles/mL) was applied to carbon-coated, glow-discharged copper grids (EMS CF400-CU-UL) and incubated for 2 minutes. Excess liquid was removed, and grids were washed three times with 1× Tris-buffered saline. Grids were then negatively stained with two sequential drops of 1% uranyl acetate (UAT) diluted in tylose water (1% UAT in deionized water (dIH2O), double filtered through a 0.22 μm filter). After drying, samples were visualized using a Hitachi 7600 TEM at 60,000× magnification and operated at 80 kV. Images were acquired with an XR80 CCD camera (AMT Imaging, Woburn, MA).

### 2.7 Single molecule localization microscopy (SMLM)

Direct stochastic optical reconstruction microscopy (DSTORM) images of EVs were obtained using a Nanoimager (ONI, Oxford Nanoimaging) and the EV Profiler kit v2 with phosphatidylserine (PS) capture. Samples were labeled with GD2-AF647, CD56-AF488, and Tetra-Trio antibodies. Calibration was done with fluorescent beads detected by illumination with 488, 561, and 640 nm lasers through a 100x oil-immersion objective lens. Successful channel mapping calibration was determined by achieving high-quality point coverage. Post-acquisition spatial and statistical analyses were conducted with CODI (alto.codi.bio), utilizing the “AI EV profiling essentials Model 1”.

### 2.8 Preparation of samples for surface proteomics

70 µL of each EV sample were incubated with 0.5 µg trypsin in 100mM triethyl ammonium bicarbonate for 1 hr at 37°C to cleave the EV surface proteins. EVs were captured onto 20 µL of magnetic bead suspension using a one-pot procedure (Affinity-based magnetic bead capture of EVs via lipid–ligand interactions, enabling direct on-bead lysis and proteomic analysis.) as described previously (Liu et al. 2023) by adding acetonitrile to a final concentration of 70%, and incubating for 10 minutes at room temperature. The cleaved surface peptides were recovered from the supernatant and dried. The isolated and dried surface peptide samples were reduced and alkylated by incubation in 10 mM TCEP and 40 mM CAA for 30 min at 37°C. The samples were further digested with Lys-C (Wako) at 1:100 (wt/wt) enzyme-to-protein ratio for 2 hr at 37°C. Trypsin was added to a final 1:50 (wt/wt) enzyme-to-protein ratio for overnight digestion at 37°C. The samples were dried in a vacuum centrifuge, desalted using Top-Tip C18 tips (Glygen) according to manufacturer’s instructions and stored at -80°C. A portion of each sample was used for peptide quantitation using Pierce Colorimetric Peptide Quantitation Kit (Thermo Fisher).

### 2.9 Preparation of samples for whole EV proteomics

30 µL of each EV sample were lysed and denatured to extract proteins using the phase-transfer surfactant (PTS) aided procedure (Iliuk et al. 2020). The proteins were reduced and alkylated by incubation in 10 mM TCEP and 40 mM CAA for 10 min at 95°C. The extracted proteins were captured onto 20 µL of magnetic beads using one-pot procedure (as described previously (Liu et al. 2023)) by adding acetonitrile to 70% final concentration and incubating for 10min at room temp. The beads were then washed three times with 70% acetonitrile and excess acetonitrile solution was allowed to dry. The proteins on beads were digested with Lys-C (Wako) at 1:100 (wt/wt) enzyme-to-protein ratio for 1 hr at 37°C in 50 µL of 50mM triethyl ammonium bicarbonate. Trypsin was added to a final 1:50 (wt/wt) enzyme-to-protein ratio for 3 hr digestion at 37°C. To recover the peptides, acetonitrile was added to each sample to a final concentration of 60% and enzymes were captured onto beads. The peptide containing supernatant was dried completely in a vacuum centrifuge, desalted and stored at -80°C. A portion of each sample was used for peptide quantitation using Pierce Colorimetric Peptide Quantitation Kit (Thermo Fisher).

### 2.10 LC-MS

The dried peptide samples were dissolved at 0.1 µg/µL concentration in 0.05% trifluoroacetic acid with 3% (vol/vol) acetonitrile. 5 μL of each sample was injected into an Ultimate 3000 nano UHPLC system (Thermo Fisher Scientific). Peptides were captured on a 2-cm Acclaim PepMap trap column and separated on a 50-cm column packed with ReproSil Saphir 1.8 μm C18 beads (Dr. Maisch GmbH). The mobile phase buffer consisted of 0.1% formic acid in ultrapure water (buffer A) with an eluting buffer of 0.1% formic acid in 80% (vol/vol) acetonitrile (buffer B) run with a linear 90-min gradient of 6–30% buffer B at flow rate of 300 nL/min. The UHPLC was coupled online with an Exploris 480 mass spectrometer (Thermo Fisher Scientific) operated in the data-dependent mode, in which a full-scan MS (from m/z 375 to 1,200 with the resolution of 60,000) was followed by MS/MS of the 30 most intense ions (15,000 resolution; normalized collision energy - 30%; normalized automatic gain control target (AGC) – 50%, maximum injection time - 30 ms; 60sec exclusion].

### 2.11 LC-MS data processing

The raw files were searched directly against The human protein atlas (Protein Atlas version 24.0) database (www.proteinatlas.org) with no redundant entries, using Byonic (Protein Metrics) and Sequest search engines loaded into Proteome Discoverer 3.1 software (Thermo Fisher Scientific). MS1 precursor mass tolerance was set at 10 ppm, and MS2 tolerance was set at 20 ppm. Search criteria included a static carbamidomethylation of cysteines (+57.0214 Da), and variable modifications of oxidation (+15.9949 Da) on methionine residues, and acetylation (+42.011 Da) at the N terminus of proteins. Search was performed with full trypsin/P digestion and allowed a maximum of two missed cleavages on the peptides analyzed from the sequence database. The false-discovery rates (FDR) of proteins and peptides were set at 0.01. All protein and peptide identifications were grouped, and any redundant entries were removed. Only unique peptides and unique master proteins were reported.

### 2.12 Label-free quantitation analysis

All data were quantified using the label-free quantitation node of Precursor Ions Quantifier through Proteome Discoverer v3.1 (Thermo Fisher Scientific) and Perseus software (version 1.6.15.0). Intensities of peptides were extracted with initial precursor mass tolerance set at 10 ppm, minimum number of isotope peaks as 2, maximum allowed retention time difference (ΔRT) of isotope pattern multiplets of 0.2 min, peptide-spectrum match (PSM) confidence FDR of 0.01, maximum RT shift of 5 min. Pairwise protein ratios were calculated as the geometric mean of log-transformed peptide-level ratios, with 100 set as the maximum allowed fold change. Hypothesis testing was done by ANOVA.

### 2.13 Comparison of surface and whole EV proteins

Proteins were classified into three groups: (1) proteins detected uniquely on the EV surface, (2) proteins detected uniquely in whole EVs, and (3) proteins found in both. A Venn diagram depicting the difference and the intersection of the EV surface/whole proteins was prepared with VennDiagram 1.7.3 R package (Chen and Boutros 2011). Additionally, we compared our results with the ExoCarta database (Keerthikumar et al. 2016), of the top 100 protein commonly identified in EVs.

### 2.14 Identification of brain-enriched and neuron-specific markers

EV surface proteins were filtered to include only those detected in all EV surface samples (stable expression/detection). The Human Protein Atlas (HPA) database (Thul and Lindskog 2018) was used to annotate these proteins with protein expression levels in human organs and mRNA expression in brain cell types. A list of primarily brain-derived markers was generated that satisfied two criteria: (1) “high” annotated protein expression in brain tissue and (2) expression in no more than three other tissues. Proteins were further subsetted for neuron specificity using mRNA expression levels (HPA) in various brain cell clusters. Neuron-specific proteins were defined as those with expression ratios <0.25 for all non-neuronal:neuronal cell cluster comparisons. Neuronal marker expression in tissues and brain cells was visualized using ggplot2 3.5.2 R package (Wickham 2016).

### 2.15 Outlier removal, differential expression analysis, and candidate biomarker identification

Outliers with z-score absolute value > 3 were removed. For missing value data, proteins detected in ≥80% samples of any one class (AD or cognitive normal) were kept (Yang et al. 2015), with no imputation of missing values (see Supplementary Figure 1 for missing value distribution). All abundances for each sample were further normalized by subtracting the median from each sample abundance and log2 transformed. Results of unpaired two-tailed Student’s t-test were controlled for FDR using the Benjamini-Hochberg method (Benjamini and Hochberg 1995). Proteins with adjusted p-value (padj) < 0.05 and absolute log2 fold change (log2FC) > 1 were considered to be significantly differentially expressed proteins (DEPs). Differential expression volcano plots were made with ggplot2 3.5.2 R package (Wickham 2016). The NeuroPro database (Askenazi et al. 2023) was used to identify DEPs that were reported in previous AD proteomic studies. DEPs identified in our study and at least five studies in the NeuroPro database and with common direction of regulation (upregulation or downregulation) were considered potential biomarkers for AD.

### 2.16 Random forest model construction

To evaluate the discriminative ability of candidate biomarkers, we constructed a random forest model based on the caret 6.0.94 (Kuhn 2008) and randomForest 4.7.1.1 R packages (Liaw and Wiener, n.d.). In the first round of the model, we used all potential AD biomarkers as feature variables and set the number of trees (’ntree’) to 200 for model training. To evaluate the performance and generalization ability of the model, we adopted five-fold cross-validation and used the pROC 1.18.5 R package (Robin et al. 2011) to construct receiver operating characteristic (ROC) curves and measured the classification performance of the model through the average area under the curve (AUC) index. Importance of each potential AD biomarker was assessed based on the mean decrease in Gini coefficient, reflecting the contribution of each feature to reducing the Gini impurity of nodes in the random forest. The five proteins with the highest importance scores were used as characteristic variables to construct the second round of the random forest model, while the remaining parameters remained unchanged.

### 2.17 Functional enrichment analysis

Gene ontology (GO) enrichment analysis of biological processes, cellular components, and molecular functions was done for AD vs CN DEPs, proteins uniquely presented on EV surface, proteins uniquely presented in whole EVs, and proteins presented in both EV surface and whole EVs using clusterProfiler 4.6.2 (Yu et al. 2012). Results were visualized with the ggplot2 3.5.2 (Wickham 2016). Interactions between proteins associated with the identified GO terms were visualized with the String database (Szklarczyk et al. 2023).

### 2.18 Confocal Airyscan microscopy of iNEVs

iNEVs were immunolabeled as previously described (Yao et al. 2025). Purified iNEVs (∼10^9^) were permeabilized with 0.001% Triton X-100 for 5min and sedimentated using 10% poly-ethylene glycol (PEG) by centrifugation at 3,500 x g for 10 min. The iNEVs pellets were resuspended in 200 µL of 1XPBS and were co-incubated with an antibody against the NEV marker VAMP2 (Vilcaes et al. 2021) and either an antibody against CNTNAP2 or NrCAM. For co-labeling of STX1B, an alternative VAMP2 antibody was used. After incubation and thorough washing, iNEVs were incubated with species-appropriate secondary antibodies conjugated to Alexa Fluor 488 or 568. All primary/secondary antibody details and dilutions are in supplementary table 4. Immunolabeled iNEVs were mounted on microscope slides with mounting medium and covered with #1.5 glass coverslips.

Immunofluorescence images were acquired using a Zeiss LSM 980 confocal microscope equipped with Airyscan, using a 100x/1.46-numerical aperture oil-immersion objective. Alexa Fluor 488 and 568 were excited using 488 nm and 561 nm lasers, respectively, with laser power set to ≤1%. All images were acquired in super-resolution (SR) configuration with an xy resolution of 1224 x 1224-pixels.

### 2.19 Single-EV flow cytometry analysis (FCA)

#### Preparation of iPSC-Derived Neuronal EVs (iNEVs) and Brain-Derived EVs (BDEVs)

Two 45-µL pools for both CN and AD-iNEVs were prepared by combining 5 µL from three technical replicates for each of three biological replicates. Brain-derived EVs (BDEVs) from post-mortem CN and AD donors were analyzed individually. For each BDEV sample, 5 µL of EV suspension was brought to 250 µL with PBS-1X and processed identically to iNEVs as detailed below.

#### Fluorescent Labeling of Total EVs

Pooled EV suspensions (45 µL) were brought to 250 µL with PBS-1X, mixed 1:1 with 250 µL of 40 µM Violet Succinimidyl Ester (VSE; Thermo Fisher Scientific, C34557), and incubated for 4 hr at 37°C in the dark. Excess dye was removed using three rounds of ultrafiltration (500 µL 100 kDa MWCO Amicon filters, Millipore,#UFC510024), yielding a final retentate of ∼100 µL. The retentate was then brought to 1.3 mL with PBS-1X and incubated with 1 µL FcR blocker (Miltenyi Biotec, #130-059-901) for 2 h at room temperature with rotation. Samples were subsequently divided into 200 µL aliquots for antibody staining.

#### Antibody Staining of Intact EVs

All staining was performed under non-permeabilized conditions to evaluate EV surface epitopes. 10 µL of the appropriate antibody solution was added to VSE-labeled EV aliquots (200 µL) to achieve a final antibody concentration of 0.2 ng/µL. Antibodies to CNTNAP2, STX1B, and NrCAM were conjugated to AF647 using Alexa Fluor® 647 Conjugation Kit (Fast) (Abcam# ab269823) kit, where as ATP1A3 was detected using appropriate Alexa Flour secondary antibodies (Supplementary table 4). Isotype controls were mouse IgG1 κ (BioLegend #400113) and mouse IgG2a κ (Invitrogen #12-4724-81). Staining was done overnight at 4°C in the dark with gentle rotation. Fc-mediated interactions were minimized by pre-incubating VSE-labeled EVs with FcR blocker (Miltenyi #130-059-901) for 2 hrs at room temperature prior to the addition of primary antibodies.

#### Single-EV FCA

Single-particle flow cytometry of EVs was performed on a CytoFLEX SRT (Beckman Coulter) using an optimized workflow adapted from methodologies described previously (Nogueras-Ortiz et al. 2024). VSE-labeled EVs were diluted immediately before acquisition to maintain event rates between 200 and 300 events per second to avoid coincident particle detection. For all analyses, the VSE+ gate defined the total EV population, and every subsequent gate was drawn within this gate. Antibody-stained intact EVs were analyzed by plotting fluorescence intensity versus violet side scatter (vSSC), used as the nanoscale sizing parameter, with size calibration (100-300 nm) established using Spherotech fluorescent nanobeads and displayed in each panel. Neuronal markers (CNTNAP2, STX1B, NrCAM) and the NEV benchmark marker ATP1A3 were quantified as the percentage of VSE⁺ EVs within gates defined by matched isotype and secondary-only controls. Dual-marker analyses employed species-specific primary and appropriate secondary antibodies for two-color detection. All samples were acquired under identical voltage, threshold, and compensation settings, with consistent VSE⁺ and antibody gate boundaries across CN and AD groups.

### 2.20 Immunoprecipitation of plasma NEVs

#### Antibody Biotinylation

Antibodies listed above were conjugated to biotin. For L1CAM, a biotinylated antibody was available (Invitrogen #13-1719-82). For antibodies in sodium azide buffers, buffer exchange was done with three rounds of ultrafiltration using 500 µL centrifugal ultrafiltration devices with a 3 kDa MWCO (Millipore #UFC500324). Samples were centrifuged at 14,000 x g until a retentate of ∼100 µL remained in additive-free PBS-1X. Protein concentration was determined using the Pierce BCA Protein Assay Kit (Thermo Scientific #23225), and antibodies were biotinylated with the EZ-Link NHS-SS-PEG4-Biotin kit (Thermo Scientific #21442), following the manufacturer’s instructions. Reactions were incubated overnight at 4°C. The following day, unreacted biotin was removed using three additional rounds of ultrafiltration (3 kDa MWCO). Final protein concentration was measured as above, biotin incorporation was confirmed using the Pierce Biotin Quantitation Kit (Thermo Scientific #28005), and biotinylated antibodies were stored at 4°C for short-term use.

#### Separation of Total Plasma EVs Using SmartSEC™

Total EVs were separated from 250 µL of human plasma using the SmartSEC™ High-Definition EV Isolation System (System Biosciences #SSS096A-1) according to the manufacturer’s instructions. The initial EV-containing flowthrough (250 µL) was combined with an additional 250 µL of EV flowthrough obtained during the column wash (500 µL total).

#### NEV IP

4.5 µg of biotinylated antibody in 50 µL IP Buffer (1% BSA, 0.1% Tween-20 in PBS 1X) were added to the 500 µL SmartSEC EV preparation and incubated at 4°C with gentle rotation. Dynabeads™ MyOne Streptavidin T1 (Invitrogen #65602) were washed three times with 1 mL of IP Buffer using a magnetic rack. After the final wash, beads were resuspended in an equal volume of IP Buffer and blocked for 30 minutes at room temperature with gentle rotation. 75 µL of pre-blocked beads were added to each EV-antibody mixture and incubated 2 hours at room temperature with gentle rotation. After incubation, beads were washed twice with 500 µL IP Buffer without mixing to minimize loss of captured EVs. The final bead pellet was briefly centrifuged in a tabletop microcentrifuge to remove residual buffer. EVs were eluted by adding 57.3 µL of 0.1 M glycine (pH ∼2.5) and incubating 10 minutes at room temperature, with vigorous vortexing at the beginning and end. Samples were briefly centrifuged to collect liquid from the tube walls and placed on a magnetic rack. Eluates were transferred to clean microtubes containing 5 µL of 1 M Tris-HCl to neutralize pH and stored at −80°C until single-molecule array (SIMOA) assays.

### 2.21 SIMOA measurements

Lysates of immunoprecipitated NEVs were prepared immediately prior to SIMOA. 24.2 µL NEVs (p-Tau217 assays) or 21 µL NEVs (p-Tau181 and p-Tau231) were lysed with 1.5X volume of 1X RIPA buffer prepared from 10X RIPA (EMD Millipore #20-188) supplemented with protease inhibitors (Thermo Scientific #78427) and phosphatase inhibitors (Thermo Scientific #1861279). Samples were frozen at −80 °C, thawed, and incubated overnight at 4 °C to ensure complete extraction of EV-associated phospho-tau. Lysates were diluted 5-fold in assay diluent prior to SIMOA measurement according to the standard protocol.

## 3 RESULTS

### 3.1 Neuronal differentiation and EV characterization

A workflow for the generation of iPSCs and neurons, separation and characterization of EVs, and data processing procedures is shown in Figure 1A. Human iPSC-derived neurons from CN and AD donors were successfully differentiated and characterized prior to EV isolation. At day 50 of differentiation, both CN and AD cultures exhibited well-developed neuronal morphology with extensive neurite networks and robust MAP2 staining, confirming neuronal identity and comparable maturation across conditions (Figure 1B). EVs separated from conditioned media were next characterized for size, concentration, morphology, and canonical surface markers based on MISEV guidelines (Welsh et al. 2024). Nano-flow cytometry analysis revealed a predominant population of small EVs with diameters primarily ranging between ∼50-150 nm diameter range in both CN and AD samples, with a peak around ∼60-80 nm (Figure 1C). While overall size distributions were similar, AD-derived EVs showed an increased particle concentration across several size bins compared to controls. Transmission electron microscopy further confirmed the presence of intact, round to cup-shaped vesicles with diameters consistent with small EVs, with no obvious morphological differences between CN and AD samples (Figure 1D). Single-molecule localization microscopy (SMLM) demonstrated the presence and nanoscale clustering of canonical EV tetraspanins CD9, CD63, and CD81 on EVs from both groups (Figure 1E). Quantitative cluster analysis indicated comparable proportions of single- and multi-tetraspanin-positive EV subpopulations in CN and AD samples, supporting the preservation of canonical EV surface signatures across conditions. Together, these data confirm successful neuronal differentiation and the isolation of well-characterized, small EVs from both CN and AD iPSC-derived neurons.

**Figure 1.**
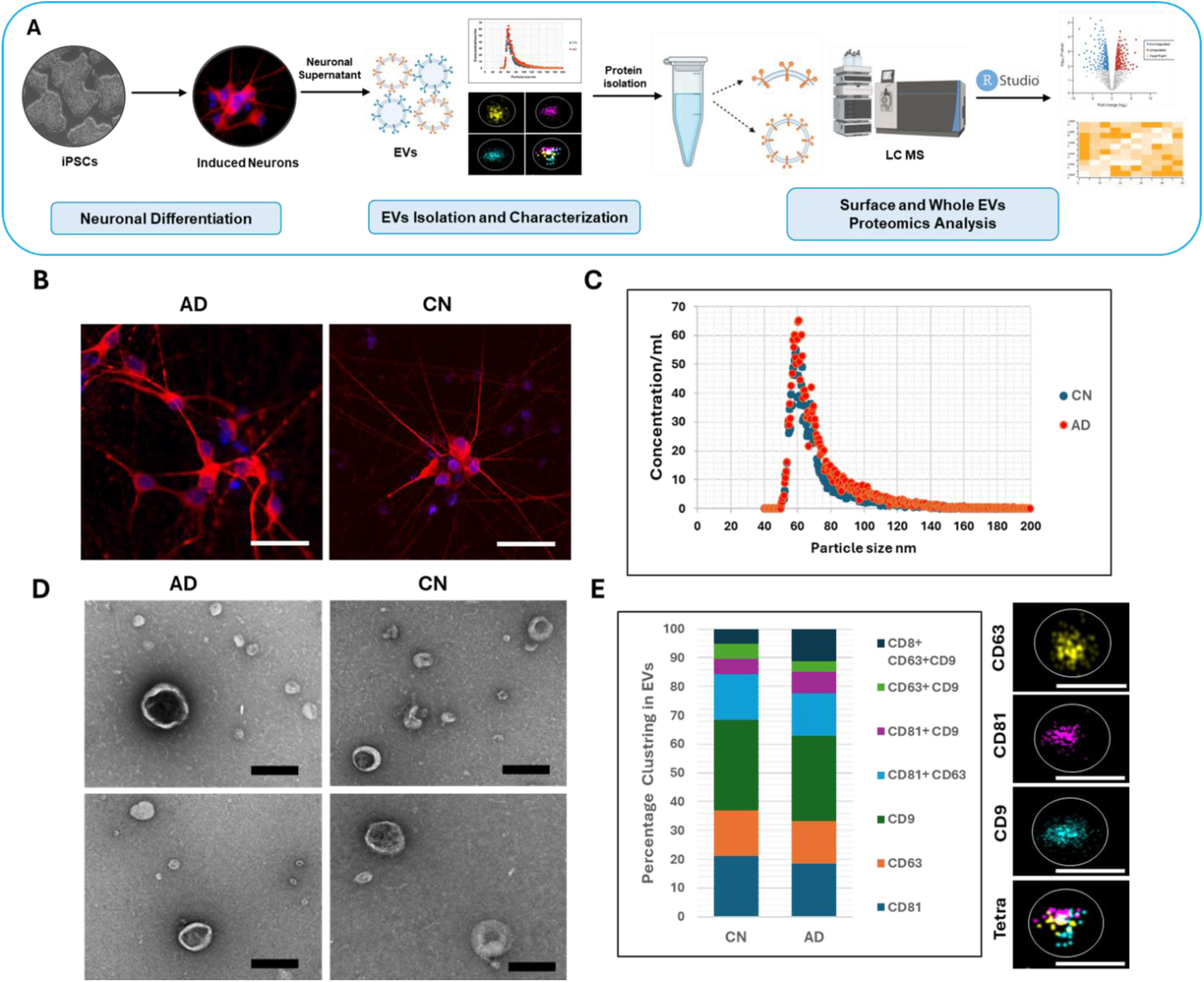
Study workflow and characterization of iPSC-derived neurons and iNEVs from CN and AD samples. (A) Experimental workflow for EV surface and whole EV protein profiling. (B) Representative immunofluorescence images of CN and AD iPSC-derived neurons at day 50 of differentiation, stained for the neuronal marker MAP2 (red) and nuclei (DAPI, blue), demonstrating comparable neuronal morphology and network formation. Scale bars= 50μm. (C)Nano-flow cytometry analysis showing particle size distribution and concentration of neuron-derived EVs from control (blue) and AD (orange) samples, presented as the average of all CN and all AD biological replicates. (D) Transmission electron microscopy images of isolated EVs from control and AD samples, revealing typical round to cup-shaped vesicles consistent with EV morphology. Scale bars = 100 nm. (E) Single-molecule localization microscopy of EVs labeled for the canonical tetraspanins CD9, CD63, and CD81, demonstrating the presence and nanoscale clustering of single- and multi-marker-positive EV subpopulations in both control and AD samples. Quantification of tetraspanin clustering is shown as a percentage distribution of EV subpopulations. Scale bars =100nm.

### 3.2 Comparative proteomic characterization of the EV surface and whole EVs

After basic characterization of EVs, as described above, we performed proteomics for 1) proteins liberated from the EV surface by proteolytic shaving and 2) whole EVs (including both surface and internal proteins). We identified a total of 5,129 proteins after data processing, including 1,360 proteins uniquely detected on the surface and 1,037 proteins identified in whole EVs (Figure 2A). Detected proteins were cross-referenced with the top 100 most commonly identified EV proteins as reported in the ExoCarta database. As shown in Figure 2B, 93% of these proteins were found in the EV surface proteome and 92% in whole EVs.

**Figure 2.**
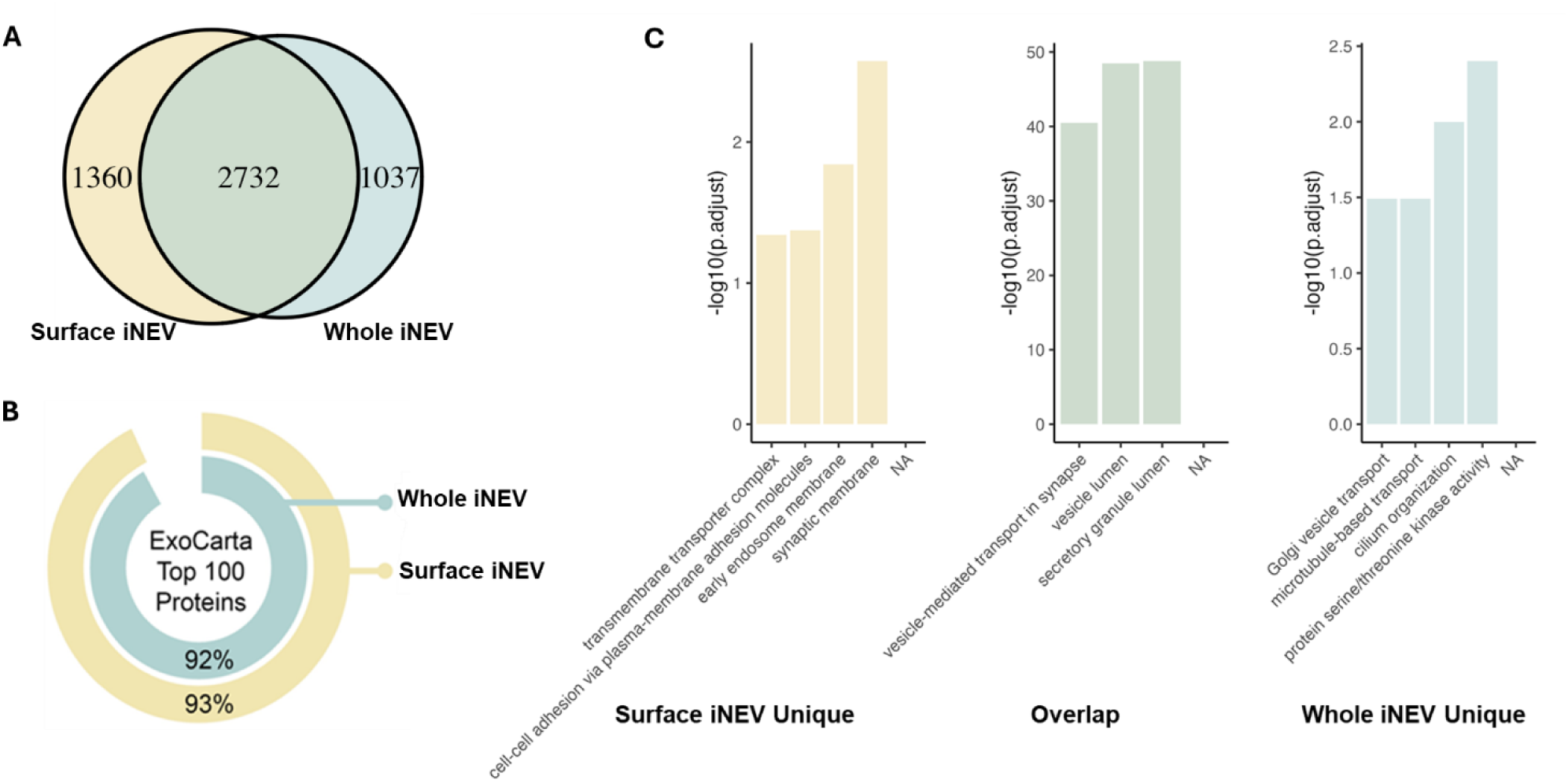
Characterization of EV Surface and Whole EV Proteomes. (A) Comparison of protein lists between EV surface and whole EV samples (B) Detection of ExoCarta top 100 commonly identified EV proteins in our EV surface and whole EV fractions. (C) GO enrichment analysis of proteins uniquely identified in the EV surface, uniquely identified in the whole EV, and shared between the two.

The EV surface and whole EV proteomes were next subjected to Gene Ontology (GO) analysis (Figure 2C). Some proteins were detected only on the EV surface but not in the whole EVs. Such surface-specific proteins may truly be distributed on the outer membrane of EVs. However, due to their relatively low abundance in whole EVs, the signals are prone to be diluted or masked by the internal high-abundance proteins, thus failing to be identified in mass spectrometry analysis. These proteins often represented GO terms such as “cell-cell adhesion via Plasma-membrane adhesion molecules”, “synaptic membrane”, and “transmembrane transporter complex” (Figure 2C).

Conversely, proteins detected only in the whole EV samples were associated with GO terms related to internal cellular processes, such as “Golgi vesicle transport”, “microtubule-based transport”, and “protein serine/threonine kinase activity” (Figure 2C). Proteins detected both on the surface and in whole EVs may be more abundant and thus more easily identified by mass spectrometry. These proteins were often associated with GO terms such as “Vesicle-mediated transport in synapse”, “regulation of trans-synaptic signaling”, and “axon development” (Figure 2C).

### 3.3 Differential expression and cross-study validation of AD-associated whole iNEV proteins

To explore the differences in the composition of EVs between AD and CN, we conducted a differential expression (DE) analysis of the whole EV proteome after outlier removal and missing value filtering. 115 significantly differentially expressed proteins (DEPs: padj < 0.05 and |log2FC| > 1) were identified (Figure 3A, B). Among which 38 were upregulated and 77 were downregulated in the AD group. Using the NeuroPro database, we identified 91 proteins that were reported to be significantly differentially regulated in AD both in this study and in others. Of these, 11 were reported by at least five other studies (Askenazi et al. 2023) and with consistent direction of regulation: 4 proteins were consistently upregulated in AD, and 7 proteins were consistently downregulated (Figure 3A, Supplementary Table 6). We defined these proteins as potential targets, as they may be used to immunocapture NEV populations that differ between AD and CN controls (Figure 3B).

**Figure 3.**
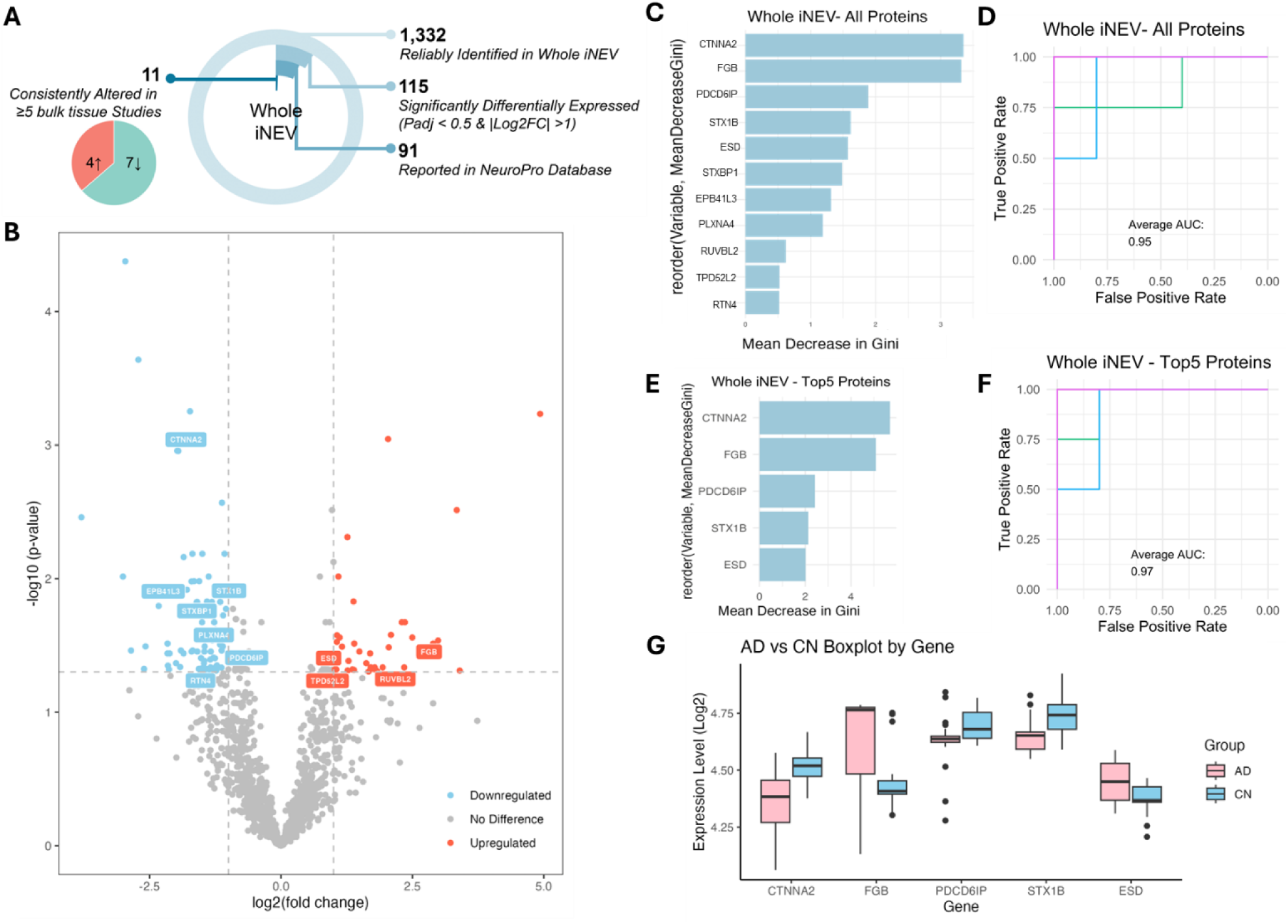
Differentially Expressed Proteins (DEPs) between AD and CN in whole iNEVs. (A)Multi-layered pie chart illustrating the number of identified DEPs. The outermost ring represents the total number of proteins detected after outlier and missing value filtering. The second ring indicates the number of significantly identified DEPs. The third ring shows the subset of DEPs that have also been reported as significantly differentially expressed between AD and healthy controls in other AD proteomics studies recorded in the NeuroPro database. The innermost ring represents DEPs that were reported in at least five NeuroPro studies with consistent dysregulation direction (upregulation/downregulation). (B) Volcano plot of DEPs. The dashed lines indicate thresholds of padj < 0.05 and |log2FC| > 1. Red dots represent significantly upregulated proteins, blue dots represent significantly downregulated proteins, and grey dots indicate proteins without significant differences. The top 11 AD-associated DEPs based on consistent reporting in the literature are labelled. (C) Importance ranking of AD-associated DEPs based on mean decrease in Gini from the machine learning model constructed with all top 11 DEPs. (D) ROC curve of the machine learning model constructed with all top 11 DEPs. (E) Importance ranking of the top 5 AD-associated DEPs based on the mean decrease in Gini from the machine learning model constructed with only these 5 DEPs. (F) ROC curve of the machine learning model constructed with the top 5 DEPs. (G) The expression levels of the top 5 DEPs in whole EVs from AD and CN groups.

To evaluate the ability of these proteins to distinguish between whole EVs from AD compared to CN, we constructed a random forest machine learning model based on this protein set. The importance ranking of each 11 proteins in the model (based on mean decrease in Gini) and the ROC curve are shown in Figures 3C and 3D. To further optimize the model, we selected the top 5 proteins based on the ranking of importance and reconstructed the model. In this simplified model, the order of importance of the five proteins remained consistent (Figure 3E), and the AUC value of the model increased by 2% to 97% (Figure 3F); thus, demonstrating that removal of lower-importance features reduces model complexity and overfitting, resulting in improved discriminatory performance of our model. The expression levels of these five proteins in the whole EV samples of AD and CN are shown in Figure 3G.

### 3.4 Differential expression and cross-study validation of AD-associated iNEV surface proteins

The same analysis was done for EV surface proteins, for which a total of 282 DEPs were identified (128 upregulated and 154 downregulated in AD) (Figure 4A, B). The NeuroPro database yielded 40 DEPs that were reported in at least five other studies (17 upregulated in AD, 23 downregulated) (Supplementary Table 5). The results of the model constructed with these 40 EV surface proteins are shown in Figures 4C and D. After selecting the top 5 EV surface AD biomarker and reconstructing the model, the AUC increased by 4% to 96% (Figure 4E, F), again demonstrating that a small, stable set of EV surface proteins is sufficient to achieve high classification accuracy, supporting their potential utility in translational applications. Expression levels are presented in Figure 4G.

**Figure 4.**
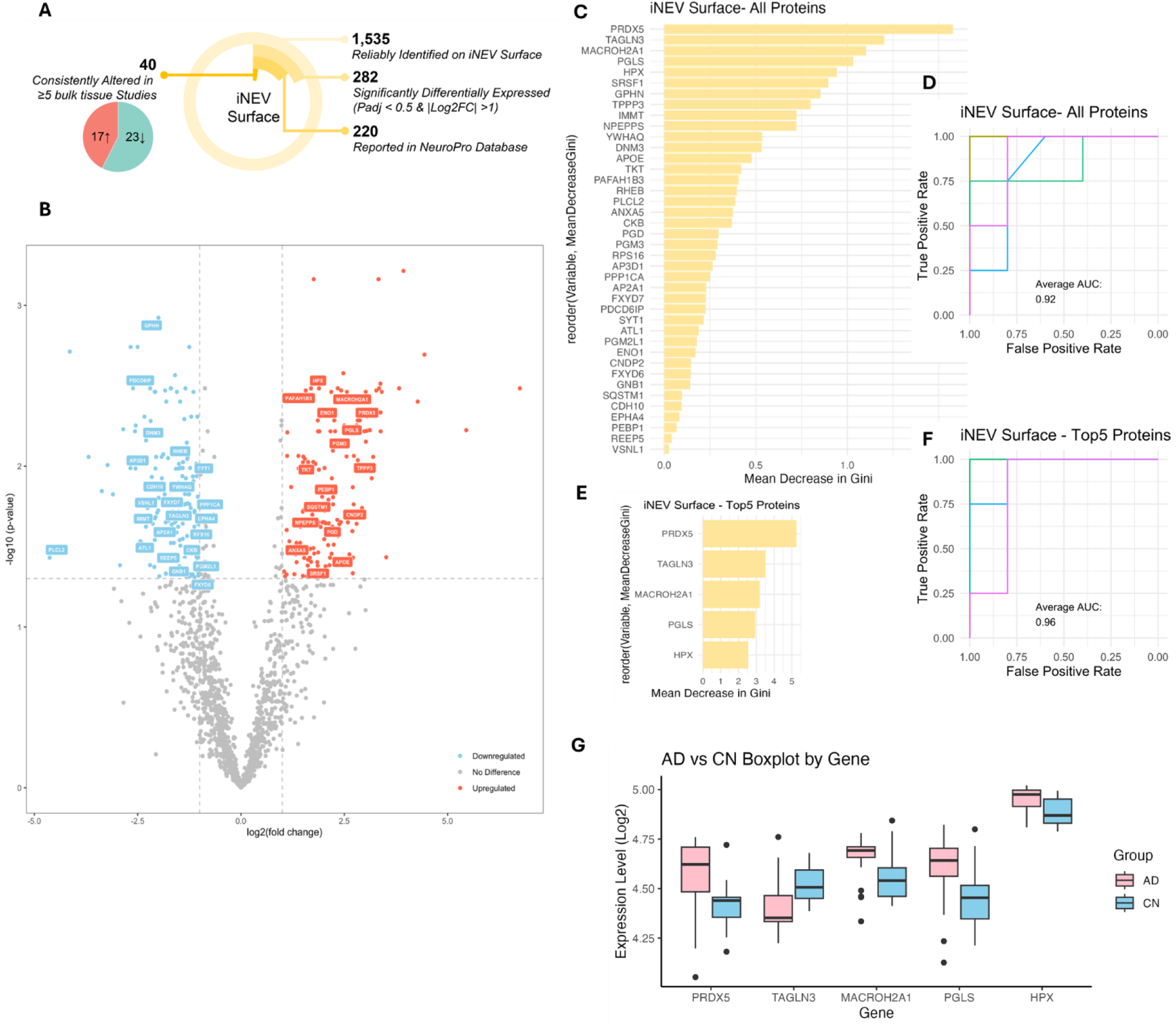
DEPs between AD and Healthy Controls in iNEVs Surfaces. (A) Multi-layered pie chart illustrating the number of identified DEPs. (B) Volcano plot of DEPs. (C) Importance ranking of all potential AD-associated DEPs based on mean decrease in Gini from the machine learning model constructed with all potential biomarkers. (D) ROC curve of the machine learning model constructed with all AD-associated DEPs. (E) Importance ranking of the top 5 AD-associated DEPs based on mean decrease in Gini from the machine learning model constructed with only the top 5 DEPs. (F) ROC curve of the machine learning model constructed with the top 5 AD-associated DEPs. (G) The expression levels of the 5 most significant DEPs in the surface of EVs from AD and CN.

Compared with whole EVs, the EV surface had more protein expression differences between AD and CN. For all DEPs identified in this study, all DEPs cross-validated with the NeuroPro database, or the subset of cross-validated DEPs that were consistently reported by at least five other studies, detected EV surface quantities were always higher than those in whole EVs (Supplementary Figure 2A). Furthermore, the log2FC distribution of DEPs on the EV surface is also broader than that of whole EVs, suggesting more sensitive detection of AD-associated proteins (Supplementary Figure 2B).

### 3.5 Functional enrichment analysis reveals AD-associated biological pathways in whole and surface EV proteomes

To better understand how EVs may be involved to AD pathology, we performed Gene Ontology (GO) enrichment analysis on the DEPs identified in both whole and surface EV proteomes. GO terms with padj < 0.05 were considered statistically significant. Figure 5A presents the top 5 enriched GO terms (with the lowest padj) across Biological Process (BP), Cellular Component (CC), and Molecular Function (MF) categories for whole EV DEPs. Several enriched GO terms were related to protein homeostasis, including “protein stabilization,” “regulation of protein stability,” “scaffold protein binding,” and “ATP-dependent protein folding chaperone,” suggesting that EVS may reflect or even mediate abnormalities in protein homeostasis in AD. Protein-protein interaction (PPI) network analysis of DEPs involved in these protein homeostasis-related GO terms revealed a tightly connected network centered around GAPDH, HSPA4, HSPD1, and CCT6A (Figure 5B). Expression levels of these DEPs in whole EVs are shown in Figure 5C.

**Figure 5.**
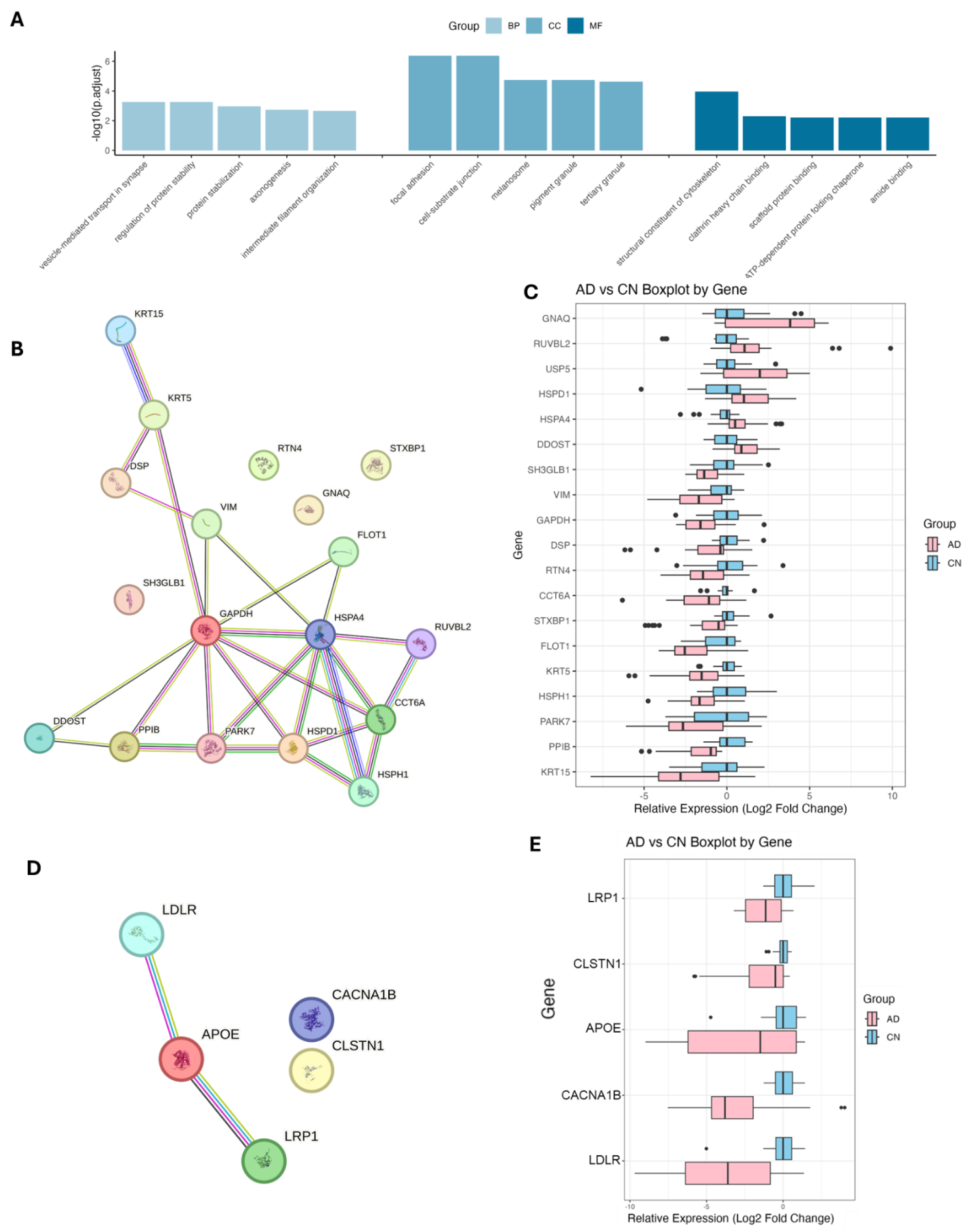
GO Enrichment Analysis of DEPs in whole EVs. (A) Top 5 enriched GO terms for BP, CC, and MF categories in whole EV DEPs. (B) PPI network of whole EV DEPs involved in protein homeostasis-related GO terms. (C) EV expression levels of whole EV DEPs involved in protein homeostasis-related GO terms. (D) PPI network of whole EV DEPs involved in the “amyloid-beta binding” GO term. (E) Full EV expression levels of whole EV DEPs involved in the “amyloid-beta binding” GO term.

Interestingly, one of the top-ranked GO terms in whole EVs was “amyloid-beta binding,” a function directly relevant to AD pathogenesis. Among the five DEPs involved in this term, LDLR, APOE, and LRP1 showed strong connections in the PPI network (Figure 5D). Moreover, these five DEPs were all significantly downregulated in AD (Figure 5E). The term “amyloid-beta clearance,” while not ranking in the top five, was statistically significant (padj = 0.008) in the BP category for whole EVs. These DEPs included LRP1, APOE, LDLR, and RAB5A (all downregulated in AD) (Supplementary Figure 3).

EV surface DEPs (Figure 6A) were associated with neuroplasticity, including “axonogenesis,” “axon development,” and “regulation of neuron projection development,” all of which may be involved in cognitive decline and neurodegeneration in AD. The PPI network of these DEPs revealed a core including PLXNA1, FYN, NRCAM, NTRK2, NCAM1, and GAP43 (Figure 6B; expression levels in 6C). Moreover, “cadherin binding” showed particularly strong enrichment (padj = 5.29e-14). In the corresponding PPI network, HSPA8, CAPZB, and PFN1 formed the core (Figure 6D; expression levels in 6E).

**Figure 6.**
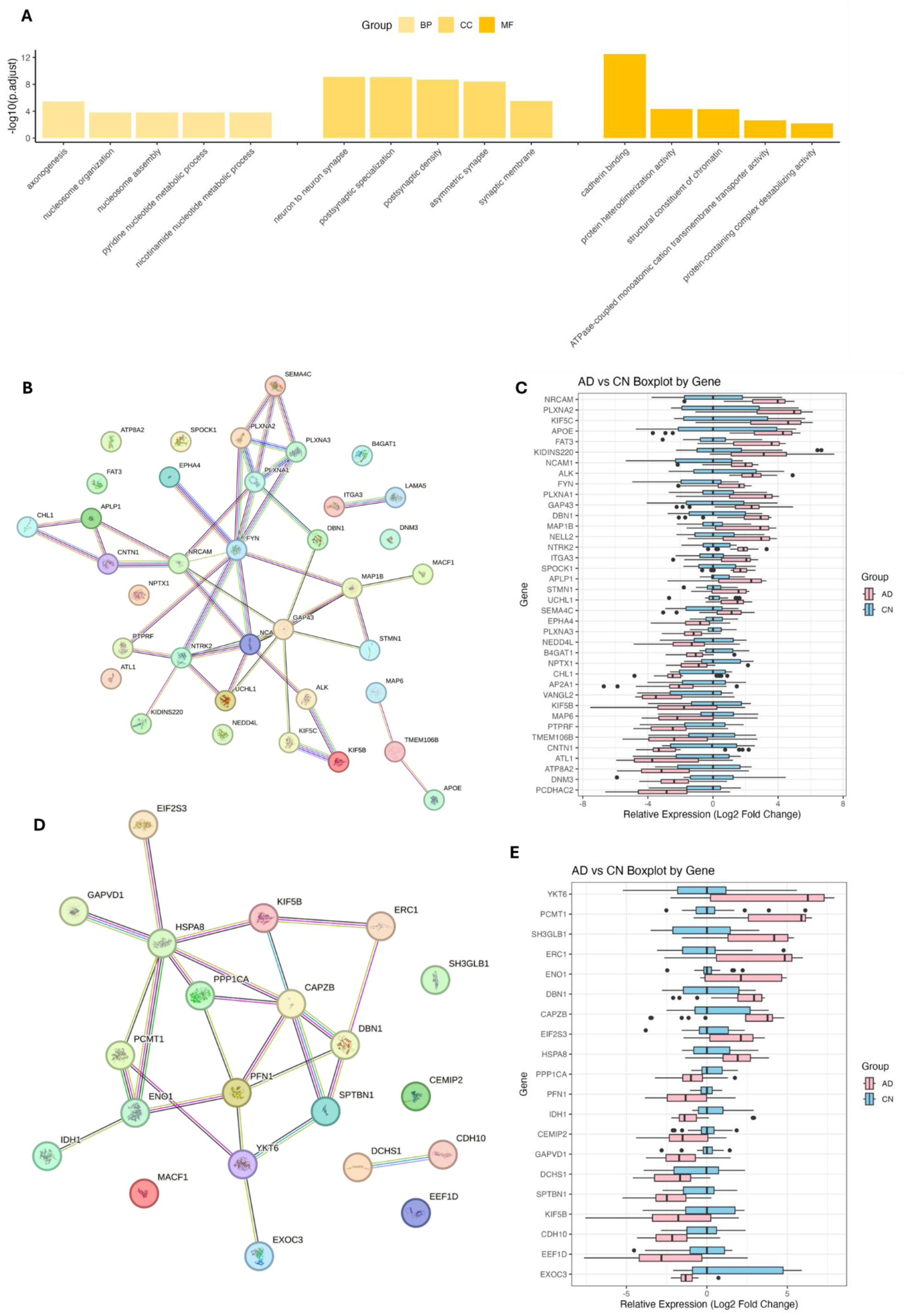
GO Enrichment Analysis of DEPs in EV surface. (A) Top 5 enriched GO terms for BP, CC, and MF categories in EV surface DEPs. (B) PPI network of EV surface DEPs involved in neuroplasticity-related GO terms. (C) EV surface expression levels of EV surface DEPs involved in neuroplasticity-related GO terms. (D) PPI network of EV surface DEPs involved in the “cadherin binding” GO term. (E) EV surface expression levels of DEPs involved in the “cadherin binding” GO term.

### 3.6 Identification of novel neuron-specific EV surface markers

To identify proteins that may be used for immunocapture of NEVs from plasma, EV surface proteins were ranked by abundance (Fig. 7A) and cross-referenced with The Human Protein Atlas to identify proteins with neuronal bias, minimal expression in blood/immune, mesenchymal, muscle, endothelial, and glial compartments (Fig. 7B), and expression across brain regions, including cerebral cortex, cerebellum, hippocampus, caudate, and amygdala, with little or no signal in non-brain tissues (Fig. 7C). Across our EV surface protein datasets, CNTNAP2 and STX1B were high-abundance, neuron-restricted surface proteins and were selected as core markers for subsequent capture of neuron-derived EVs from plasma and BDEVs. In contrast, ATP1A3 and NrCAM, which were previously proposed as an iNEV biomarker (Huang et al. 2023; You et al. 2023), showed appreciable expression in non-brain tissues and non-neuronal cell types (Fig. 7D), indicating limited specificity for NEVs. Collectively, these analyses support the investigation of CNTNAP2 and STX1B for neuronal EV enrichment.

**Figure 7.**
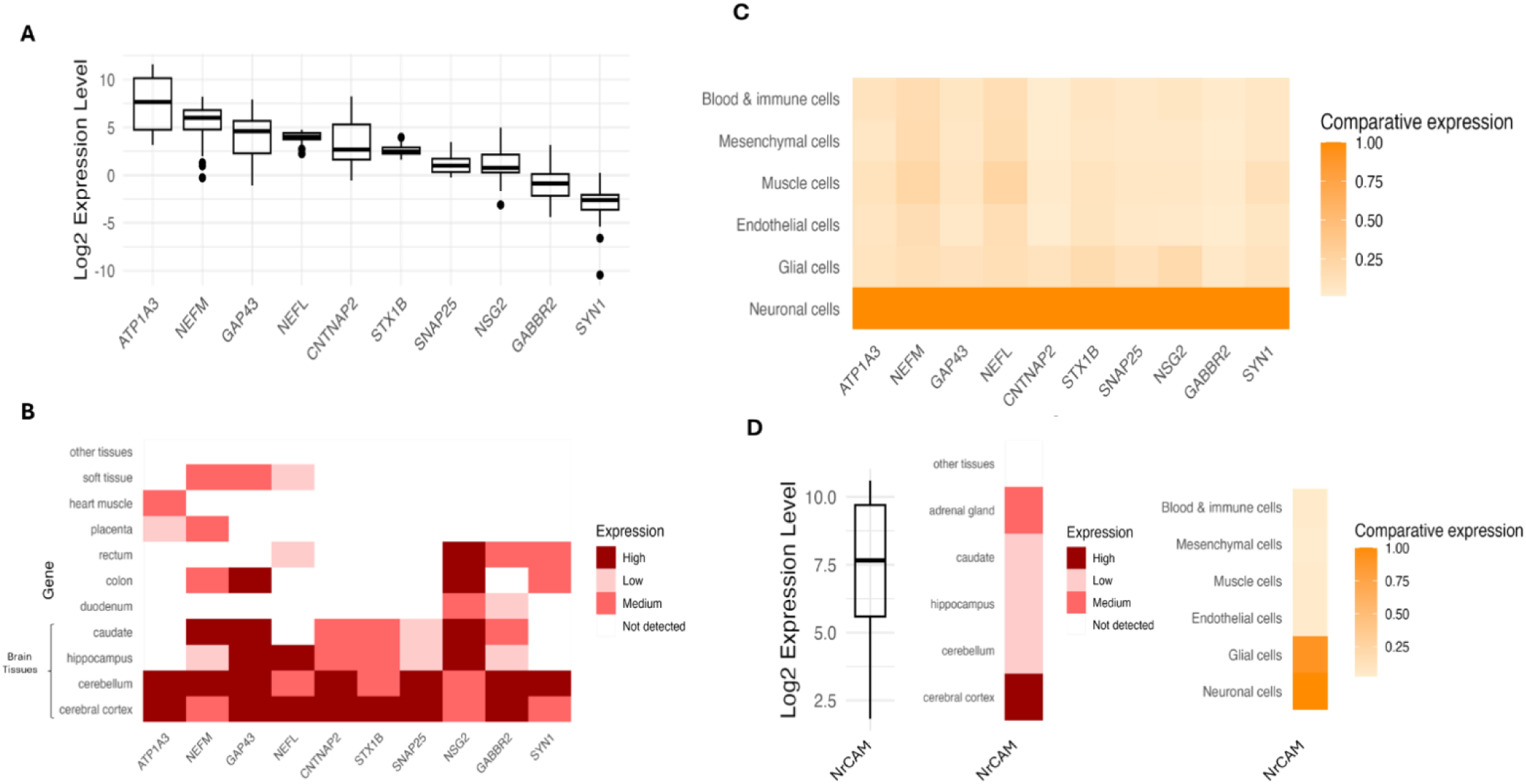
Identification of highly expressed neuronal EV surface markers across all samples. (A) Ranked abundance of candidate proteins detected on the surface of plasma EVs in the EV surface–enriched fraction; candidates are ordered high to low by normalized signal. (B) Cell-type specificity of the same candidates based on Human Protein Atlas (HPA) annotations, showing expression in neurons, blood/immune, mesenchymal, muscle, endothelial, and glial cells. (C) Tissue/brain-region specificity from HPA, demonstrating enrichment across cerebral cortex, cerebellum, hippocampus, caudate, and amygdala (D) NrCAM expression profile across tissues and cell types from HPA, indicating detectable expression outside the brain and in non-neuronal cells. In all heatmaps, color intensity reflects relative expression within each dataset.

### 3.7 Validation of newly identified neuronal enrichment markers

To validate proteomic hits identifying putative neuronal EV surface proteins, we examined CNTNAP2 and STX1B on individual iNEVs and assessed their colocalization with the previously reported neuronal marker NrCAM by confocal microscopy; all three proteins localized to EVs and showed clear colocalization with the canonical EV marker VAMP2. (Vilcaes et al. 2021), confirming their presence on individual EVs (Figure 8A). Single-particle flow cytometry analysis (FCA) (Nogueras-Ortiz et al. 2024; Shen et al. 2018), was done next. EVs were labeled generically with Violet Succinimidyl Ester (VSE) (Figure 8B). Within the VSE+ population (Figure 8 C-H), CNTNAP2, STX1B, and NrCAM were detected on iNEVs (Figure 8 F-H), comparable to the previously shown neuronal EV marker ATP1A3 (Figure 8E) and consistent across EVs from AD and CN iPSC-derived neurons (Figure 8I). Assessment of double positives revealed that STX1B displayed the highest degree of overlap with ATP1A3, followed by NrCAM and CNTNAP2 (Figure 8 J-K). We also examined brain-derived EVs (BDEVs) separated from AD and CN brain tissue. All three candidate proteins were detected with similar distributions across AD and CN (Figure 8L). Together, these results corroborate the proteomic identification of CNTNAP2 and STX1B as bona fide neuronal surface proteins on EVs and support their utility as surface-accessible targets for positive selection of neuron-derived EVs in human biofluids and biomarker assessment.

**Figure 8.**
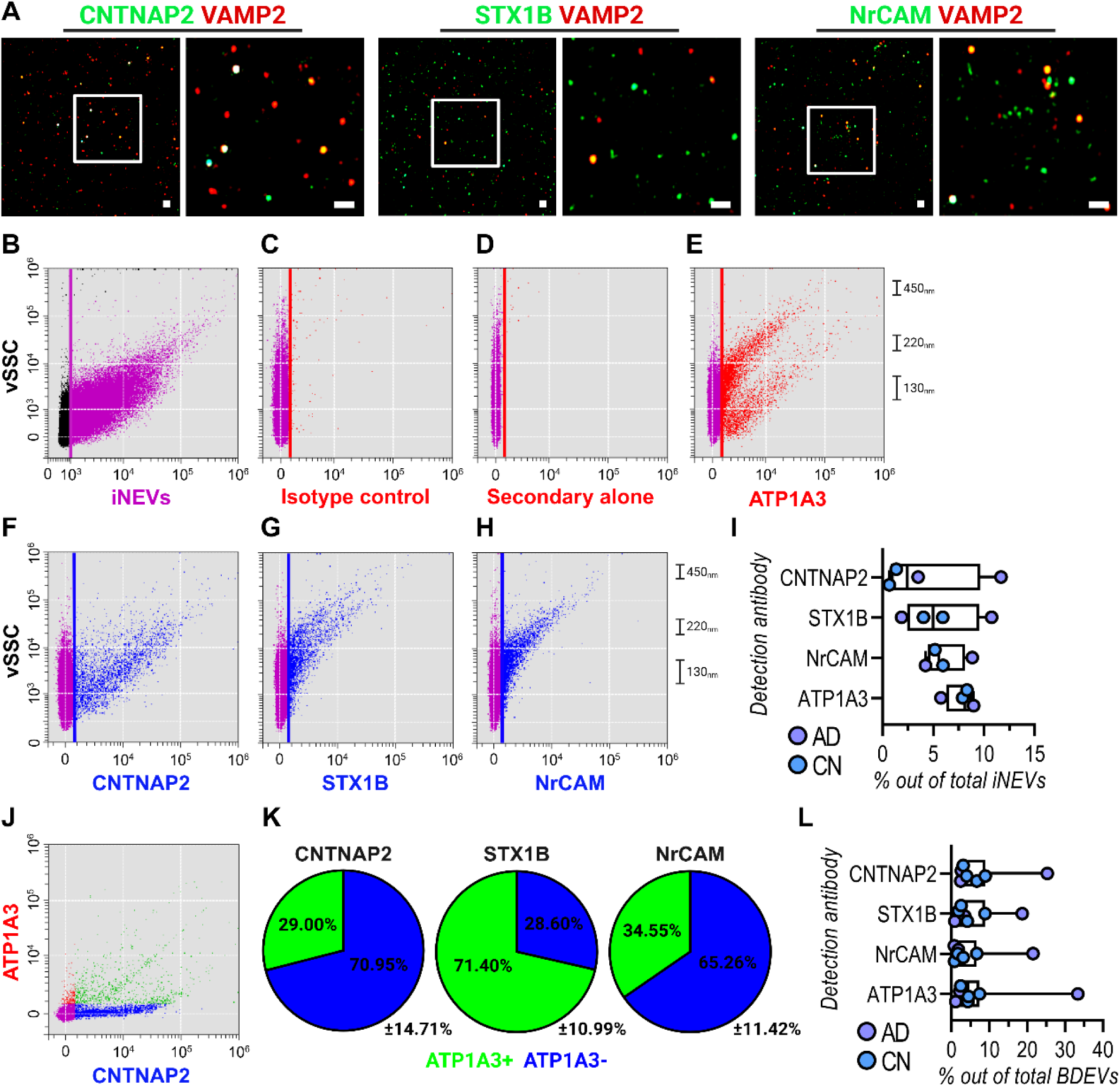
Validation of candidate neuronal markers on human iNEVs and BDEVs. (A) Confocal fluorescence microscopy of single iNEVs showing colocalization of neuronal membrane proteins CNTNAP2, STX1B, and NrCAM (green) with the canonical NEV marker VAMP2 (red). Insets show magnified fields illustrating co-localized signals in orange. Scale bars, 1 µm. (B-H) Single-EV FCA of intact iNEVs labeled with Violet Succinimidyl Ester (VSE) to define total EV events. All gating for negative controls (red) and target membrane proteins (blue) was performed strictly within the VSE+ EV population (B; magenta). Panels show VSE+ total EV gating (B), isotype control (C), secondary-only control (D), and the neuronal EV benchmark ATP1A3 (E), followed by representative FCA detection of CNTNAP2 (F), STX1B (G), and NrCAM (H). Each FCA plot displays antibody fluorescence intensity on the x-axis and violet side scatter (vSSC)-a representation for vesicle size-on the y-axis. A nanoscale reference legend is included to the right of each FCA panel, providing approximate EV size estimates (in nanometers) derived from the vSSC signal of fluorescent nanobeads. (I) Quantification of the percentage of VSE+ events double positive for each marker (CNTNAP2, STX1B, NrCAM, ATP1A3) in AD and CN iNEVs. Each data point represents the mean signal of a pooled iNEV sample generated from the conditioned media of three independent iPSC-derived neuronal lines. Box-and-whisker plots display the median, interquartile range (IQR), and whiskers extending to the full data range. (J) Representative bivariate FCA density plot illustrating double-positive iNEVs for ATP1A3 (y-axis) and CNTNAP2 (x-axis). Red and blue clusters denote single-positive ATP1A3 or CNTNAP2 events, respectively, and green events denote ATP1A3/CNTNAP2 double-positive iNEVs. (K) Summary pie charts depicting the proportion of ATP1A3+ (green) and ATP1A3- (blue) iNEVs within CNTNAP2, STX1B, and NrCAM-positive EV populations (mean ± SEM; n = 4 pooled CN and AD samples). These distributions illustrate the degree of overlap between each candidate surface marker and the established neuronal EV marker ATP1A3. (L) Box-and-whisker plots (median, IQR, and Tukey whiskers) summarizing BDEV surface-marker expression in CN and AD brain tissue. Each data point reflects the mean fluorescence signal from two technical replicates for each biological sample (n = 3-4 per group).

### 3.8 Discrimination of AD and CN by CNTNAP2 immunoseparation

We next examined whether these novel neuronal surface markers could precipitate EVs enriched in AD-relevant phosphorylated tau cargo from human plasma. We first evaluated phosphorylated tau levels in NEV immunoprecipitates (IPs) derived from SEC-isolated total EVs from CN older adults. These pre-IP samples served as the baseline against which enrichment was assessed. IPs using antibodies against CNTNAP2, STX1B, NrCAM, and ATP1A3 markedly increased detectable p-Tau217 and p-Tau181 relative to matched total EV inputs (Figure 9A), demonstrating that all four neuronal surface proteins could precipitate EVs carrying phosphorylated tau. L1CAM, another neuronal EV marker previously used by our group for immunoprecipitation of neuronal EVs, was included as an additional comparator.

**Figure 9.**
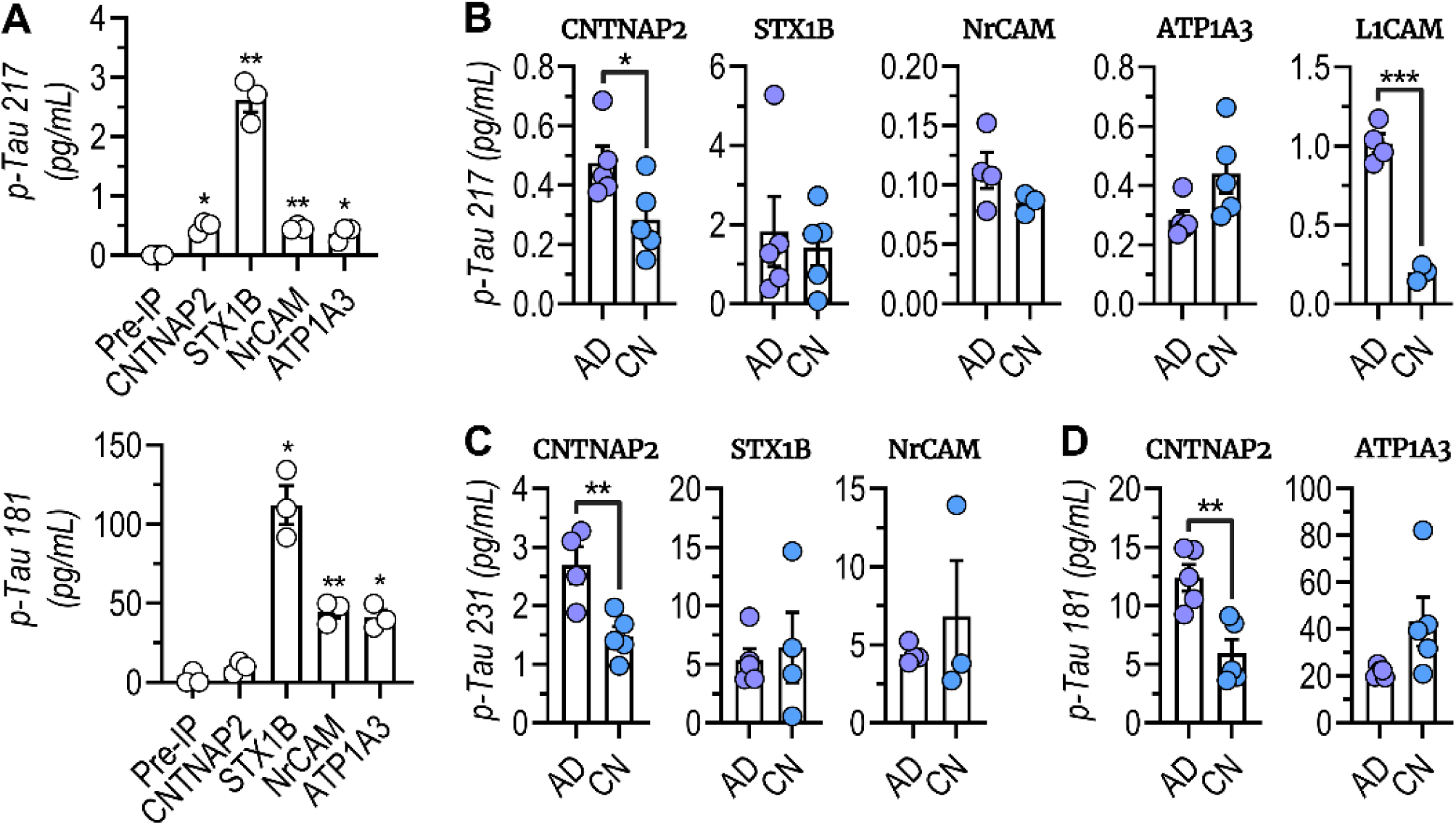
SIMOA quantification of phosphorylated tau species in NEV immunoprecipitates from human plasma. (A) Enrichment of phosphorylated tau species following NEV immunoprecipitation (NEV IP) from plasma of cognitively normal older adults (n = 3; mean age = 77 ± 4.9 years, all male). Plasma was subjected to IP using CNTNAP2, STX1B, NrCAM, or ATP1A3 antibodies, and SIMOA was used to quantify p-Tau217 (top) and p-Tau181 (bottom). Pre-IP plasma levels are shown for comparison. All four neuronal surface-marker IPs selectively enriched phosphorylated tau relative to input plasma. (B-D) SIMOA quantification of phosphorylated tau species in NEV IPs from plasma of individuals with AD and CN older adults. The AD and CN groups each consisted of 5 female participants (ages 68-83 years; AD mean age = 76.4 ± 4.5, CN mean age = 77.2 ± 5.5). EVs were immunoprecipitated using antibodies against CNTNAP2, STX1B, NrCAM, ATP1A3, or L1CAM, and phosphorylated tau was quantified by SIMOA. Panel B shows p-Tau217, panel C shows p-Tau231, and panel D shows p- Tau181 levels in NEVs from AD and CN participants recovered via IP targeting the markers indicated. Individual data points represent plasma donors; bars denote group means ± SEM. Occasional bars with smaller sample numbers reflect limited aliquot availability for certain markers; in these cases, leading candidate markers and essential comparators were prioritized for analysis. Comparisons in (A) were analyzed using a repeated-measures one-way ANOVA across matched SEC-total EV and corresponding NEV IP conditions, whereas all AD-CN comparisons in the remaining panels were assessed using unpaired, parametric two-tailed t-tests.

We next compared NEV IPs from AD and CN across three phospho-tau biomarkers quantified by SIMOA. For p-Tau217, CNTNAP2-IP NEVs exhibited a clear and statistically robust elevation in AD relative to CN (Figure 9B), mirroring the AD-associated increase also recovered by L1CAM IP. In contrast, ATP1A3-IP NEVs showed a modest decrease in p-Tau217 in AD, an inverted directionality consistent with the behavior of ATP1A3-positive NEVs as previously reported (You et al. 2023). Analysis of p-Tau231 (Figure9C) and p-Tau181 (Figure 9D) demonstrated the same overall patterns: CNTNAP2-IP discriminated AD and CN, STX1B-IP and NrCAM-IP yielded minimal or no differences, and ATP1A3-IP showed the opposite trend.

## 4 DISCUSSION

This study set out to address a central bottleneck in the field of liquid biopsies for brain disorders: the specific and reliable separation of NEVs from human plasma. EVs provide a window into the state of their cells of origin and are increasingly recognized as a rich source of biomarkers across diseases (Yuana et al. 2013; Tran et al. 2025). However, the complexity of plasma and the overlapping expression of many proteins across tissues have made it difficult to attribute circulating EVs to specific organs.

Our proteomics-first approach was designed to overcome these limitations by nominating and validating proteins that are brain-enriched, EV-associated, and detected in most of our iNEV samples (with a strict “80% rule” for the downstream inclusion). Several markers we identify improve the signal-to-noise ratio for neuronal cargo, defined here as consistent detection of neuronal EV proteins across samples relative to sporadic or background EV proteins compared to the previously identified neuronal markers. Consistent with the idea that surface selection shapes the observed cargo, we also observed marker-dependent differences in EV content, suggesting that different epitopes enrich partially distinct NEV subpopulations (which may be attributable to different neuronal subtype, maturation state, or subcellular origin).

Differential expression analysis of proteins reliably detected in whole and surface iNEV proteomes showed large differences between AD and CN. Most DEPs detected in whole and surface datasets were also reported in the NeuroPro database, emphasizing reproducibility. GO analysis of the iNEV DEGs shows that they are enriched in pathways related to protein homeostasis and neuroplasticity, which is consistent with Gomes’ et.al. report that EVs can be involved in AD pathology through “protein degradation” and “synaptic plasticity and homeostasis” (P. Gomes et al. 2022). Importantly, we found that DEPs of whole iNEVs are also enriched in amyloid-related pathways, which strongly suggests a potential involvement of EVs in the regulation of amyloid-beta dynamics in AD pathogenesis. Thus, further supporting that iNEVs are related to AD pathology and indicating that iNEVs may also influence the AD progression by regulating the accumulation and clearance of the amyloid plaques (Figure 5). The relative downregulation of beta-amyloid-binding proteins on iNEVs of AD origin (LDLR, APOE, and LRP1) suggest that these NEVs may be less able to capture and help clear beta-amyloid from the extracellular environment, potentially prolonging its toxic presence. Furthermore, the high significance of the “cadherin binding” GO term suggests that the surface of EVs may play a role in AD pathogenesis through cadherin-related pathways (Figure 6). Our further analysis showed that the machine learning models based on the AD-associated DEPs identified in whole iNEVs (CTNNA2, FGB, PDCD6IP, STX1B, ESD) and on iNEV surface (PRDX5, TAGLN3, MACROH2A1, PGLS, HPX) can distinguish between AD and control neurons of origin (AUC = 0.97 and 0.96). All of these support our hypothesis that the iNEVs offer great potential for AD prediction and diagnosis.

To identify iNEV surface proteins that might be useful to separate neuronal EVs from the much larger background of heterogeneously sourced EVs in blood plasma, we used The Human Protein Atlas to assess protein expression by tissue and RNA expression by brain cell type for the proteins identified in most of our iNEV surface samples. Only those proteins that were specifically and highly expressed in brain tissue and neuronal cells were retained. 10 proteins satisfied our expression thresholding criteria: ATP1A3, NEFM, GAP43, NEFL, CNTNAP2, STX1B, SNAP25, NSG2, GABBR2, and SYN1 (Figure 7). We identified 2 proteins (CNTNAP2 and STX1B) with the highest potential as candidates for further validation based on their strong brain and neuronal enrichment, consistent detection across samples under the ≥80% inclusion criterion, and predicted accessibility on the EV surface. Since, ATP1A3, L1CAM and NrCAM are popular markers for NEV immunocapture, we also included them in various downstream experiments for comparison. Confocal microscopy and high-resolution FCA confirmed that CNTNAP2, STX1B along with previously known marker NrCAM and ATP1A3 reside on the surface of intact NEVs (Figure 9), providing the necessary spatial context for their use as immunocapture targets in biofluids. These data established CNTNAP2 and STX1B proteins as bona fide, surface-accessible membrane markers on NEVs and demonstrate their presence across both iNEVs and BDEVs.

Our SIMOA results (Figure 9), however, can be interpreted as challenging the prevailing assumption that the diagnostic utility of an NEV surface target is dictated primarily by its neuronal specificity: that is, a marker proven to be neuron-specific is presumed to be an effective handle for enriching EVs of neuronal origin and for measuring AD or other brain disease-relevant biomarkers (Manolopoulos et al. 2025). Our findings partially revise this assumption, suggesting important additional parameters: differences among NEV subpopulations in cargo loading and directionality of biomarker differences. By comparing IP of multiple neuronal surface proteins, we show that different NEV membrane targets may enrich for different NEV subpopulations, resulting in profoundly different biomarker profiles for AD (and, potentially, other brain diseases).

CNTNAP2 IP consistently enriched for phospho-tau species relevant to AD, particularly p-Tau217, whereas STX1B and NrCAM IPs showed negligible AD-CN separation. Most strikingly, ATP1A3, a rapidly adopted neuronal EV benchmark, showed an inverse AD signature, with lower phospho-tau in AD than in CN. This pattern is consistent with previous observations (You et al. 2023) and indicates that ATP1A3 may isolate an NEV population that is fundamentally different from the one identified in earlier studies of NEVs by NrCAM and L1CAM IPs (Fiandaca et al. 2015; Kapogiannis et al. 2019; Eren et al. 2020). In contrast, L1CAM, one of the longest-used neuronal EV targets, displayed the expected AD-associated increase in phospho-tau, further validating its ability to access AD-relevant NEV pools (Nogueras-Ortiz et al. 2024; D. E. Gomes and Witwer 2022). Together, these findings demonstrate that neuronal specificity alone is insufficient to predict whether a surface protein will provide meaningful biomarker signatures.

We propose a biologically realistic model: plasma, and probably brain tissue itself, contain multiple NEV pools that are separable by their membrane signatures, likely reflecting different neuronal subtypes, brain regional origins, proximity to the blood-brain barrier, or differential derivation from neuronal parts (soma, dendrites, neurite, synapses). Our bivariate FCA findings reinforce this model, since CNTNAP2, STX1B, and NrCAM co-localized only partially with ATP1A3, rather than marking a single, uniform NEV population (Figure 8K). Plasma NEVs thus include multiple sub-pools defined by surface phenotype, and, as our SIMOA results show, each may betray AD pathology in a different way or not at all. Under this model, CNTNAP2 and ATP1A3 identify largely non-overlapping NEV populations, and only the CNTNAP2+ EV pool is similar to the L1CAM+ pool in AD-associated phospho-tau cargo. However, CNTNAP2 displays highly brain-restricted expression, supporting its potential as a more specific brain-derived EV marker (Rodenas-Cuadrado et al. 2014; Thul and Lindskog 2018). This biological diversity has implications for biomarker development. The above-mentioned characteristics of CNTNAP2 position it as a next-generation NEV marker suited for diagnostic biomarker development, therapeutic monitoring, and, potentially, mechanistic studies. In contrast, STX1B, NrCAM, and ATP1A3, while being robust markers of neuronal identity, appear to define EV populations that are less reflective of tau pathology in plasma.

In summary, our findings reveal a previously unrecognized molecular diversity among circulating NEVs, demonstrate that the utility of NEVs as a source of biomarkers cannot be inferred solely from their neuronal specificity, and identify CNTNAP2 as a high-value target for future plasma EV diagnostics as well as mechanistic studies in AD.

## ACKNOWLEDGEMENTS

The authors would like to acknowledge our patients across North America who provided clinical data and blood samples.

## FUNDING

The study funders had no role in the study design; in the collection, analysis or interpretation of data; in the writing of the report; or in the decision to submit the paper for publication.

V.M. – NIH (1RF1AG083801); The Richman Family Precision Medicine Center of Excellence in Alzheimer’s Disease at Johns Hopkins University.

D.K. – This research was supported in part by the Intramural Research Program of the National Institutes of Health (NIH). The contributions of the NIH author are considered Works of the United States Government. The findings and conclusions presented in this paper are those of the author and do not necessarily reflect the views of the NIH or the U.S. Department of Health and Human Services.

C.G.L – This work was supported by the Richman Family Precision Medicine Center of Excellence in Alzheimer’s Disease at Johns Hopkins including significant contributions from the Richman Family Foundation, the Rick Sharp Alzheimer’s Foundation, the Sharp Family Foundation and others. Dr. Lyketsos was also supported by P30AG066507 to the Johns Hopkins ADRC

K.W.W. – Paul G. Allen Frontiers Foundation; The Richman Family Precision Medicine Center of Excellence in Alzheimer’s Disease at Johns Hopkins University

## CONSENT STATEMENT

Induced Human iPSCs used in this project have been generated from peripheral blood monocytes obtained upon fully informed consent from human subjects as outlined by the JHMI eIRB-approved research study application (NA_00045104) and consent process. All experiments were conducted according to regulations governing the use of human stem cell lines in biomedical research.

## AUTHOR CONTRIBUTIONS

V.M. conceptualized the study. W.A. and R.S. differentiated iPSC lines into neurons. W.A. and R.S. isolated extracellular vesicles. A.I., on behalf of Tymora Analytical Operations, prepared EV for proteomics, processed LC-MS/MS data, and performed label-free quantitation analysis. W.A and D.D. performed data processing, batch effect correction, and normalization, performed bioinformatics, data analysis, and figure generation using R and Biorender.com. W.A. and C.N.O. validated the findings in plasma and BdEV samples. W.A., D.D C.N.O., R.J.B. wrote and assembled the manuscript. W.A., C.N.O., R.S., D.D., R.J.B, A.I., C.G.L., K.W.W., D.K., and V.M. contributed to scientific discussion and edits.

## DATA AVAILABILITY

The data supporting the findings of this study are available from the corresponding author upon reasonable request

