## Supplementary figures for "Proteomic identification and validation of novel neuronal EV-based markers for Alzheimer’s disease biomarker discovery"

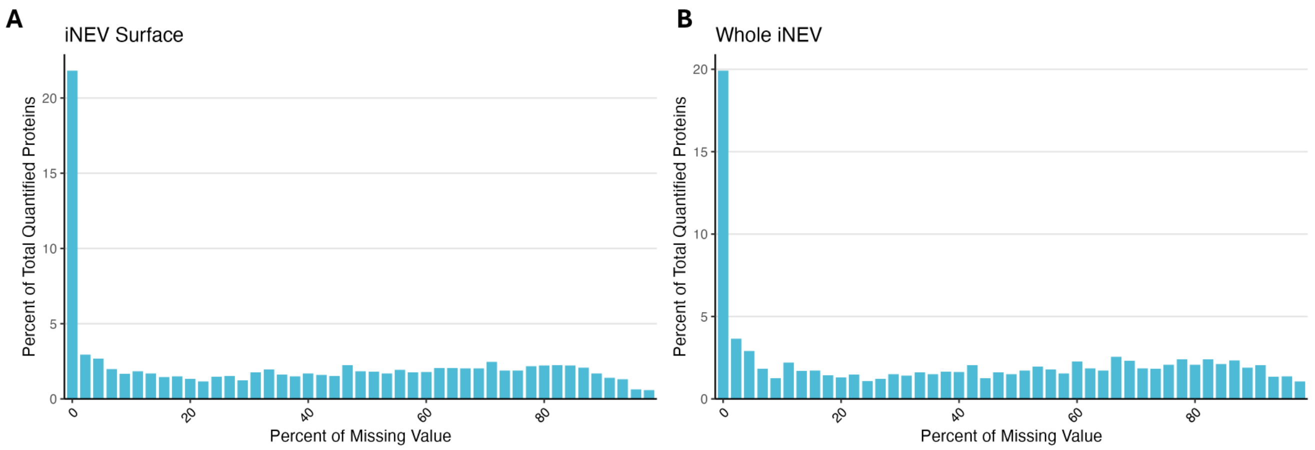


**Supplementary Figure 1. Missing value distribution of EV surface and whole EV proteomes data. a.** Missing values in EV surface data. **b.** Missing values in full EV data.

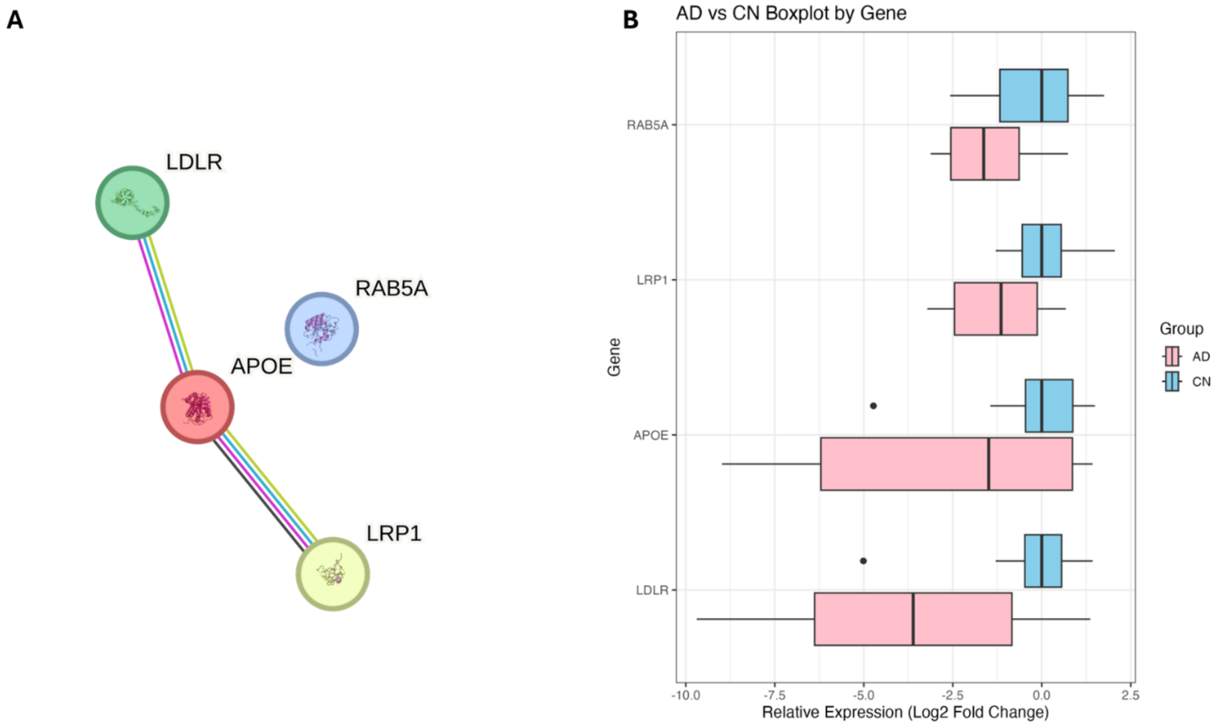


**Supplementary Figure 2. Whole EV DEPs Involved in Amyloid-beta Clearance GO Term. a.** PPI network of these DEPs. **e.** Full EV expression levels of these DEPs.

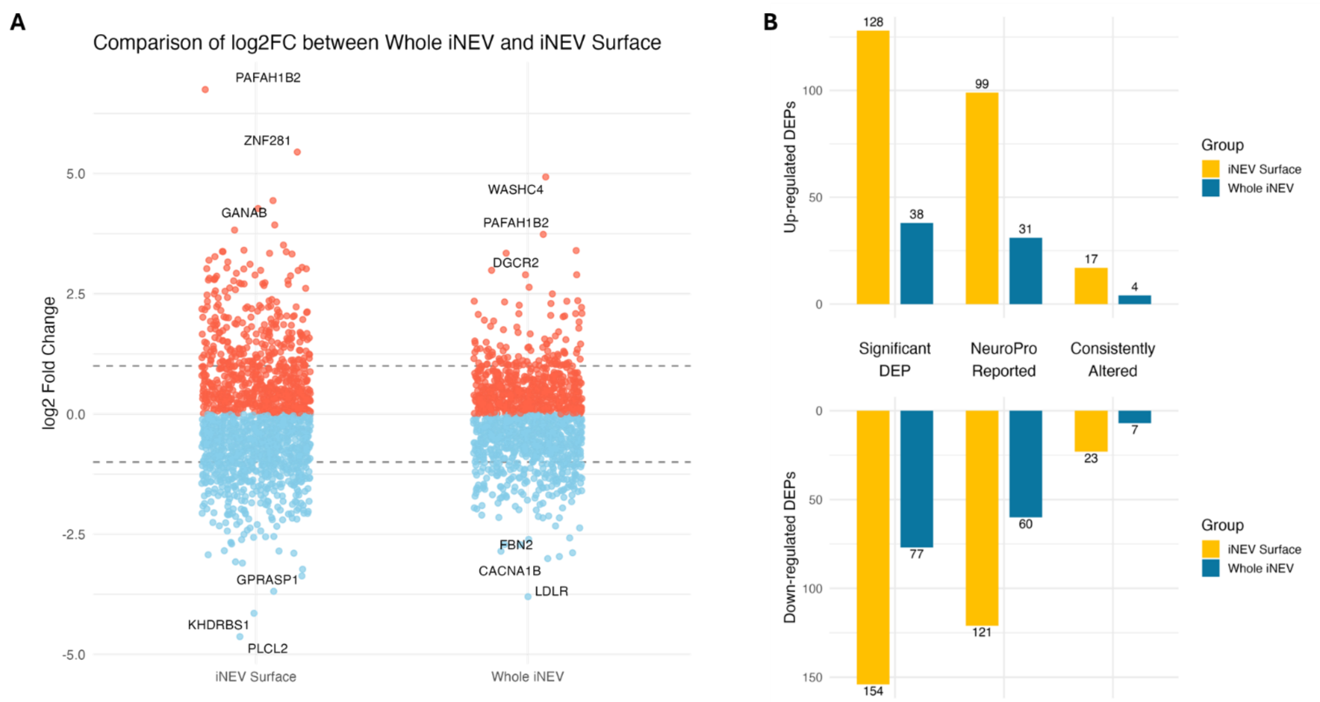


**Supplementary Figure 3. Comparison between DE Results of EV Surface and Full EV Data. a.** Log2 fold change distribution of DE results. The three proteins with the highest log2FC and the three proteins with the lowest log2FC have been labeled. **b.** DEP numbers of EV surface and full EV DE results.
